# Low-affinity binding motif in microtubule plus-end condensates specializes microtubule function

**DOI:** 10.64898/2026.06.15.732316

**Authors:** Madhurima Choudhury, Federico Uliana, Tarik Grubić, Mateusz P. Czub, Ana-Maria Farcas, Michel O. Steinmetz, Yves Barral

## Abstract

The microtubule plus-end tracking proteins (+TIPs) CLIP-170/Bik1 and EB/Bim1 form a condensate, the +TIP body, at the plus-end of most microtubules in vivo. Remarkably, however, these +TIP bodies typically impart different dynamics and interaction profiles to distinct microtubules, according to their cellular function. The molecular mechanisms underlying the functional versatility of the +TIP body are unknown. Here, we show that the +TIP Kar9 utilizes repeats of a lysine-aspartate-lysine (KDK)-centered short linear motif (SLiM) to interact with Bik1 on a restricted subset of cytoplasmic microtubules during yeast mitosis. Furthermore, these multivalent Kar9-Bik1 interactions tune the material behavior of the +TIP body to specify proper microtubule function. Indicating that KDK serves as generic Bik1-interaction motif, similar motifs are also present in Kip2, where they mediate Bik1-dependent recruitment of Kip2 to the +TIP body. Together, our study provides insights into how low-affinity Bik1 interactors diversify microtubule function by locally specializing the content and behavior of +TIP bodies.

## Introduction

How biomolecular, nanometer scale interactions drive the self-organization of cellular structures at the meso- and micrometer scales remains a key question of cell biology. Over the years, two fundamental mechanisms have taken center stage, namely the non-covalent polymerization of cytoskeletal filaments, such as actin filaments and microtubules (reviewed in (Dominguez and Holmes, 2011; Goodson and Jonasson, 2018), and the assembly of biomolecular condensates by associative, liquid-liquid phase separation (Banani *et al*., 2017). Whereas cytoskeletal filaments rely on stoichiometric, stable interactions, phase separation is driven by networks of multivalent, low-affinity and transient interactions. Remarkably, these two distinct mechanisms of organization appear to be intimately articulated with each other within cells, as an increasing number of condensates are found to associate with the cytoskeleton (Yaguchi and Woodruff, 2026). However, our understanding of how these interactions contribute to shaping cellular architecture is still in its infancy.

Indeed, the idea that the interplay of the cytoskeleton and specific condensates is pivotal for cellular architecture is supported by a growing list of examples. For instance, cell-cell contacts in tissues are largely shaped by the reciprocal reinforcement of actin filaments and membrane-associated condensates, such as N-WASP condensates at the cell cortex and ZO1 condensates at tight junctions (Wiegand et al., 2025; Beutel et al., 2019). Likewise, the nucleation and organization of spindle microtubules rely on the embedding of their minus ends in the condensate formed by the pericentriolar material (PCM) in centrosomes (Woodruff *et al*., 2017).

Here, we focus on +TIP bodies, a type of condensate that decorate the plus-ends of microtubules and which is composed of microtubule plus-end tracking proteins (Morelli et al., 2025; Song et al., 2023; Maan *et al*., 2023; Meier *et al*., 2023; Miesch *et al*., 2023). Recent studies have shown that these +TIP bodies can recruit tubulin dimers, influencing microtubule growth rates and catastrophe frequency (Miesch *et al*., 2023), collect cargoes (Chen et al., 2023; Maan et al., 2023; Chen et al., 2019), or link microtubule ends to the actin cytoskeleton (Hwang *et al*., 2003). These +TIP bodies arise from co-condensation of two classes of +TIPs: ubiquitous +TIP proteins that localize to virtually all microtubule plus-ends, and patterning +TIPs (Akhmanova and Steinmetz, 2008, 2015; Kumar et al., 2021), present on specific microtubules and lending them functional versatility. Ubiquitous +TIPs, principally the cytoplasmic linker protein (CLIPs, such as CLIP-170, Bik1 in budding yeast) and the end- binding proteins (EBs/Bim1), are highly conserved in evolution, whereas patterning +TIPs, such as the proteins MACF, APC, and SLAIN in mammals and Kar9 in budding yeast, are structurally more variable and less conserved across species. Therefore, the interplay between ubiquitous and patterning +TIPs emerges as a mechanism for specifying the behavior and cellular role of individual microtubules. However, what limits the localization of individual patterning +TIPs to destined microtubules is unclear, as are the mechanisms by which they then modulate the behavior of these microtubules.

Over the years, budding yeast has revealed to be a powerful model system for studying +TIP bodies. In these cells, the Kar9 +TIP body forms at the tip of cytoplasmic microtubules, exclusively. There, it recruits an actin-dependent motor protein, the type V myosin Myo2 (Yin et al., 2000; Kumar et al., 2021; Morelli et al., 2025), and links microtubule ends with actin filaments (Hwang *et al*., 2003). Thereby, it steers the movements of the nucleus within the cell. Strikingly, in pre-anaphase cells, the Kar9 body is restricted to the microtubules emanating from only one spindle pole body (SPB, centrosome equivalent in yeast), the old SPB, which the cell has inherited from the previous mitosis (Liakopoulos *et al*., 2003; Hotz *et al*., 2012). This ensures that the mitotic spindle, which assembles between the old and a newly synthesized SPB, is pulled along actin cables towards the mother-bud neck of the cell, i.e., the future plane of cell cleavage (Beach et al., 2000; Yin et al., 2000; Miller, Matheos and Rose, 1999). In addition, this aligns the spindle with the mother-bud axis and orients the old SPB towards the bud and hence the future daughter cell (Pereira *et al*., 2000; Pereira *et al*., 2001; Hotz *et al*., 2012). At anaphase onset, the Kar9 body harvests the forces generated by microtubule shrinkage at the bud cortex to pull half of the spindle into the bud (Morelli *et al*., 2025). Thus, this system provides an excellent case for studying how patterning +TIPs are targeted to specific microtubules and modulate their behavior.

Our recent studies have suggested that a key element of Kar9 function is to modulate the material properties of the +TIP body (Meier *et al*., 2023; Morelli *et al*., 2025). For example, multivalent Kar9-Kar9 interactions are needed for harvesting the force generated by microtubule shrinkage, probably by tuning the viscoelastic properties of the Kar9 body. Consequently, mutations affecting Kar9’s ability to phase separate impair its localization to only one side of the spindle prior to anaphase. This observation has suggested that the ability of the Kar9 body to coarsen through Oswald ripening drives Kar9’s symmetry breaking in mitosis (Meier, Steinmetz and Barral, 2024).

To further investigate how the material properties of the Kar9 body contribute to its localization and function, we dissect here how Kar9 interacts with one of the ubiquitous components of the Kar9 body, CLIP-170/Bik1. Indeed, previous studies established that the function of the Kar9 body strongly relies on multivalent Kar9-Kar9 interactions (Meier *et al*., 2023; Morelli *et al*., 2025). In addition, previous studies emphasized the importance of the ubiquitous +TIP Bik1 in determining the function of the +TIP body (Meier et al., 2023; Miller et al., 2006; Moore, Silva and Miller, 2006). However, how the behavior of ubiquitous +TIP components changes and adapts according to the function of specific +TIP bodies, and how patterning +TIPs contribute to this process is unclear. Therefore, we went on to dissect how Kar9 interacts with Bik1 and how these interactions influence the behavior and function of the entire +TIP body.

## Results

### 1. Bik1 condensates selectively enrich Kar9 in vitro

Kar9 and Bik1 interact with each other, as demonstrated by pull-down and yeast two-hybrid experiments (Moore, Silva and Miller, 2006). However, the exact modalities of this interaction is unknown. Protein-protein interactions within condensates are often of low-affinity and transient but multivalent (Banani *et al*., 2017; Shin and Brangwynne, 2017). Although methodologies to probe stable protein-protein interactions are widely established, we lack undemanding approaches to probe low-affinity interactions.

To dissect the interaction between Kar9 and Bik1, we developed a partitioning-based methodology. A series of bacterial lysates, each expressing a different Kar9 fragment C-terminally tagged with mNeonGreen (mNG) (Figure 1A, 1B, fragment 1-9) were mixed with recombinantly produced and purified Bik1 protein, and induced to phase separate by decreasing the ionic strength (from 500 to 250 mM NaCl) (Meier et al., 2023; Czub et al., 2025). The fluorescence intensities of the Kar9-tagged fragment within the Bik1 droplets and in the dilute phase were then measured by confocal microscopy. The ratio of these fluorescence intensities (K) defines the partition coefficient of a chosen fragment into the Bik1 droplet under our experimental conditions. The partitioning of mNG itself into Bik1 droplets (K=1.91 ±0.07) served as reference.

**Figure 1:**
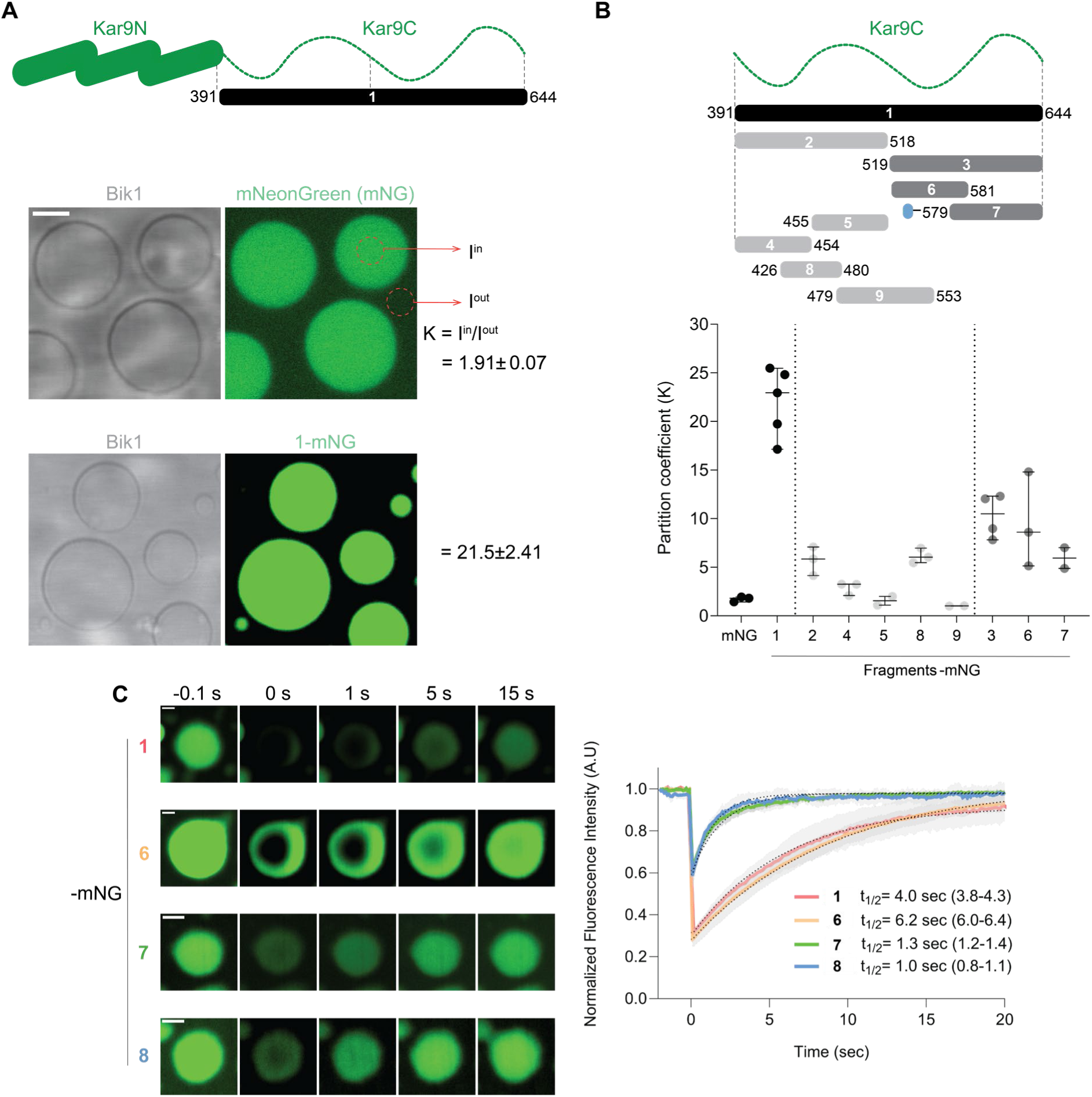
Partition of Kar9 fragments into Bik1 droplets. **Figure 1A:** Partitioning behavior of Kar9 disordered C-terminal domain (Kar9C, fragment 1) with C-terminal mNeonGreen (mNG) tag from *E. coli* BL21(RIL) lysates into Bik1 droplets (33 µM) with respect to mNG alone. Protein schematic representation shows different domains of Kar9, with Kar9N as the folded domain and Kar9C as the disordered C-terminal domain. Residue numbers are mentioned for the respective fragment in the representation. K represents partition coefficient, which is the ratio of the mean intensity within a selected region of interest in the condensed phase (I^in^) to a region of interest in dilute phase (I^out^) and is calculated from one independent experiment. Scale bar: 2 µm. Each experiment was repeated at least 2 times. Presence of full-length fragments were verified by Western Blot (Figure S1). **1B:** Partitioning behavior represented by partition coefficients of diverse fragments of Kar9 disordered C-terminal domain C-terminally tagged with mNG. Fragment 7 has an 8-amino acid long S/G linker in N-terminus (represented in blue) to aid solubility. Kar9C is the Kar9 disordered C-terminal domain. Residue numbers are mentioned for the respective fragment in the representation. Each data point represents median partition coefficient of respective fragments from at least 25 droplets in independent experiments. Bars represent median values with standard deviations. Presence of full-length fragments were verified by Western Blot (Figure S1). **1C:** Fluorescence Recovery After Photobleaching (FRAP) of fragments 1, 6, 7, and 8 (in all cases, 3 droplets were bleached within single experiments). Colored lines show mean recovery curves of three photobleaching experiments, and the shaded regions show standard deviations. Dotted lines show fitted recovery curves with 1-phase association model in GraphPad Prism 4.0. *t_1/2_* represents half recovery times with 95% confidence intervals in parentheses.

Interestingly, the disordered C-terminal domain of Kar9 (aa 391-644, fragment 1) tagged with mNeonGreen partitioned robustly into the Bik1 droplets, indicating that it contains at least one Bik1-binding interface, as suggested by previous data (Moore, Silva and Miller, 2006). To dissect which part of the Kar9 C-terminal domain interacts with Bik1, we apportioned it further into 8 fragments of different sizes (Figure 1B). Strikingly, 5 of these 8 fragments partitioned to Bik1 droplet above the level of mNG control, namely fragments 2 (aa 391-518), 3 (aa 519-644), 6 (aa 519-581), 7 (aa 579-644) and 8 (aa 426-480) (Figure 1B). Thus, the Kar9 C-terminal domain may interact with Bik1 through several independent sites. Supporting this notion, the partition coefficients of fragments 2, 3, 6, 7, and 8 are lower than that of fragment 1, consistent with multivalency increasing the interaction avidity of the Kar9 C-terminal domain for Bik1. Importantly, detection of these interactions in the crowded milieu of the bacterial lysate supports the idea that these interactions may be functional in the yeast cellular environment.

To characterize the dynamics of these interactions, we next turned to fluorescence recovery after photobleaching (FRAP). The fluorescence of Kar9 fragments was photobleached in a small area of Bik1 droplets and fluorescence recovery was monitored over time. All tested fragments recovered within seconds, indicating that interactions were short-lived and that partitioning equilibrium was rapidly reached (Figure 1C). Furthermore, fluorescence of the full-length C-terminal domain (fragment 1) and fragment 6 recovered the slowest (*t_1/2_* = 4.0 sec, 95% confidence interval: 3.8-4.3 and 6.2 sec - 6.0-6.4 -, respectively), followed by fragments 7 and 8 (*t_1/2_* = 1.3 sec - 1.2-1.4 - and 1.0 sec - 0.8-1.1-). In comparison, the fluorescence of fragment 9 fully recovered within the time of a single frame imaging (0.1 sec), indicating that its recovery time is below 0.05 sec. Thus, the recovery time of each fragment correlated generally well with its partitioning strength (Figure 1B, 1C).

### 2. Individual charged patches contribute to Kar9 C-terminal domain partitioning into Bik1 droplets

To precisely identify the molecular determinants of Kar9 C-terminal domain interaction with Bik1, we trimmed fragment 6 and 7 sequentially. Interestingly, two smaller and contiguous fragments, fragment 6.1 (amino acid 534-566) and 6.2 (amino acid 564-594) contained similar patches of alternatively charged amino acids, namely, 562-KDK, and 579-KDK (hereon named as P1 and P2 respectively) (Figure 2A). Analysis of manually aligned Kar9 sequences across fragment 6.1 and 6.2, and several fungal species showed that the Kar9 homologs also contained consistently two K/R-D/E-K/R patches, although their exact positions were not necessarily conserved across species (Figure S2B). Importantly, while the charged amino acids were the most conserved part of these alignments, its context also showed some level of conservation, with the frequent presence of hydrophobic (mainly I,L,V) as well as serine/threonine at conserved positions before and after P1 and P2. Thus, we speculated that P1 and P2 may be part of a motif that mediates Kar9 interactions with Bik1.

**Figure 2:**
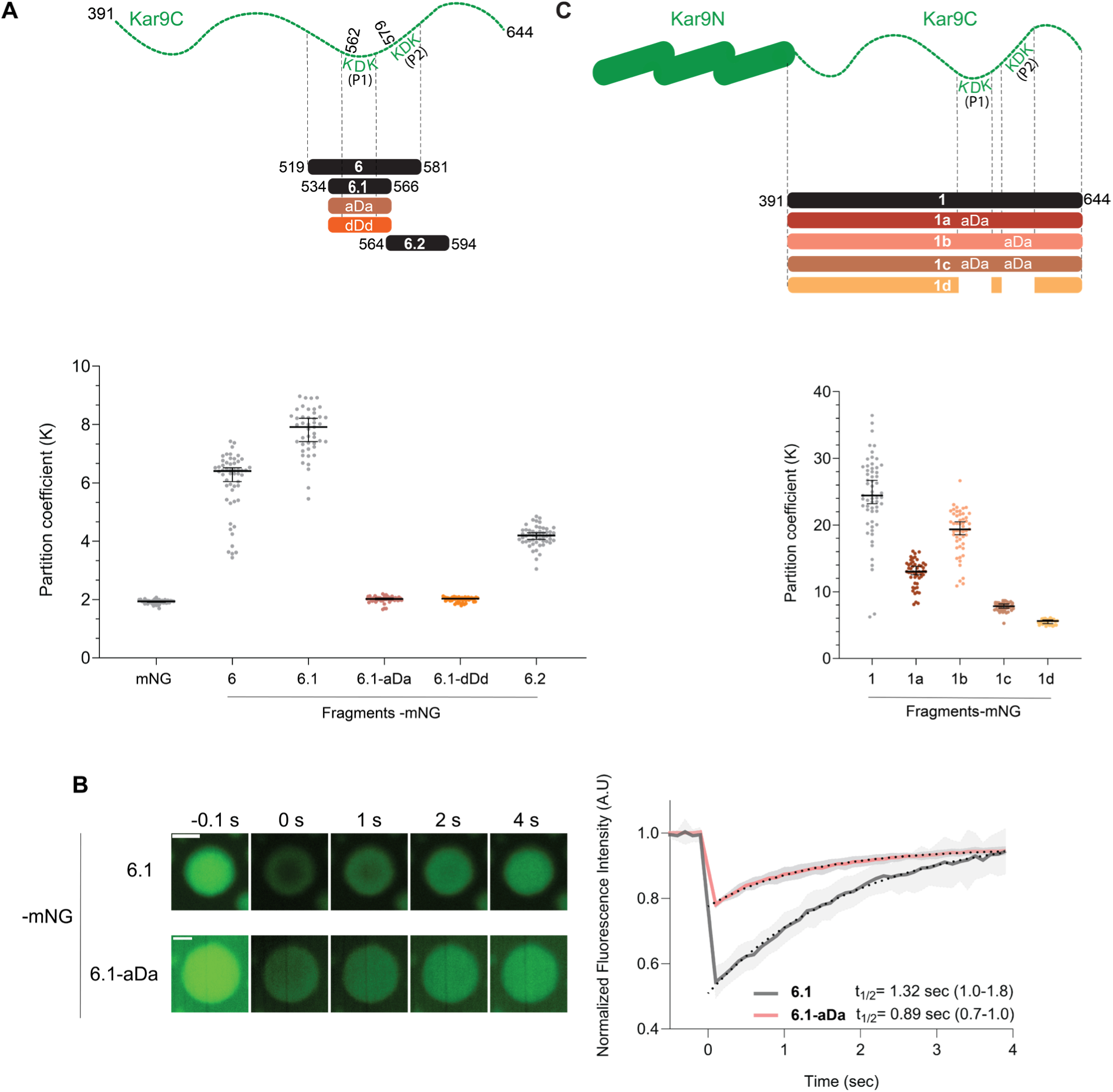
Identification of Kar9-Bik1 interaction site in vitro. **2A:** Partitioning behavior of diverse fragment 6 variants represented by partition coefficients. Kar9C is the Kar9 disordered C-terminal domain. Residue numbers are mentioned for the respective fragment in the representation. Each data point represents partition coefficient within a droplet obtained in single experiment. Each experiment was repeated at least 2 times. Presence of full-length fragments were verified by Western Blot (Figure S2A). **2B:** Fluorescence Recovery After Photobleaching (FRAP) of fragment 6.1 and 6.1-aDa (in all cases, 3 droplets were bleached within single experiments). Colored lines show mean recovery curves of three photobleaching experiments, and the shaded regions show standard deviations. Dotted lines show fitted recovery curves with 1-phase association model in GraphPad Prism 4.0. *t_1/2_* represents half recovery times with 95% confidence intervals in parentheses. **2C:** Partitioning behavior of Kar9 disordered region (fragment 1) and its mutant variants (fragment 1a, 1b, 1c and 1d) represented by partition coefficients. Protein schematic representation shows different domains of Kar9, with Kar9N as the folded domain and Kar9C as the disordered C-terminal domain. Residue numbers are mentioned for the respective fragment in the representation. Each datapoint represents partition coefficients within a droplet obtained in single experiments. Bars represent median values with standard deviations. Presence of full-length fragments were verified by Western Blot (Figure S2A).

Supporting this hypothesis, mutating the positively charged residues in P1 to alanines or aspartic acids (i.e., 562-aDa/562-dDd) abrogated the partitioning of fragment 6.1 into Bik1 droplets (Figure 2A) and substantially reduced its recovery time upon photobleaching (Figure 2B). While the expression of mutated fragment 6.2 was insufficient to test its partitioning, we were able to produce the full-length C-terminal domains mutated in P1 and P2 individually or together (mutant fragments 1a, 1b, 1c, and 1d; Figure 2C). Separately mutating each of the patches by substituting the lysine with alanine residues (fragment 1a and 1b) reduced the fragment 1 partitioning into Bik1 droplets compared to the wild type fragment (Figure 2C). Mutating or removing P1 and P2 together (fragment 1c and 1d) further lowered the partition coefficient (6<K<8, Figure 2C) of fragment 1, indicating that both patches of charged residues contribute to Kar9 interaction with Bik1, independently of each other. The fact that mutating both patches does not fully reduce the partitioning of Kar9 C-terminal domain to mNG control levels (Figure 2C, 1B) indicates that additional interaction sites are still at play, probably those present in fragment 2 (Figure 1B).

### 3. Patches P1 and P2 in Kar9 directly interact with Bik1

Our partitioning data suggest that at least two sequence patches in the C-terminal domain of Kar9 contribute to Kar9-Bik1 interaction. To determine whether P1 and P2 indeed interact with Bik1 directly, we performed chemical crosslinking mass spectrometry (XL-MS) [PMID: 26654279] on the mixture of Bim1, Bik1 and Kar9 at equimolar ratio, under dilute and condensed conditions [PMID: 39885130, PMID: 34487489]. In this approach, the nucleophilic residue lysine in proteins are covalently linked with a homobifunctional molecule (disuccinimidyl sulfoxide, DSS) in which the functional groups are separated by a rigid arm, thereby constraining linked residues within ∼30Å. After crosslinking, proteins samples were subjected to bottom-up proteolysis with trypsin or Glu-C, and the peptides were identified by mass spectrometry (crosslinked peptides identified and filtered are reported in the Supplementary Table 1) (Figure 3A, 3B, S3C, S3D).

**Figure 3:**
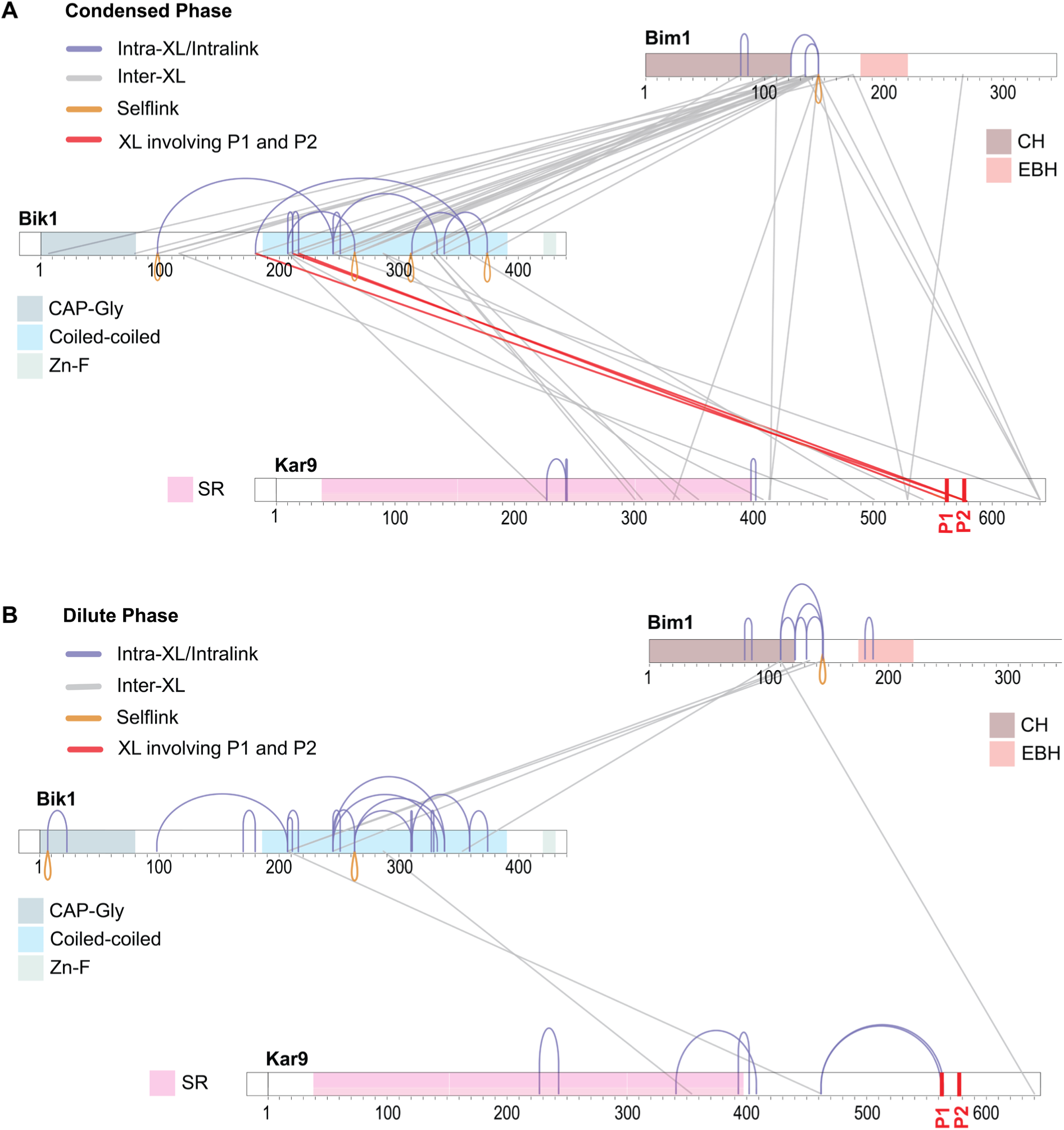
Crosslinking Mass Spectrometry of Kar9, Bik1, and Bim1. **3A and 3B:** Visualization of cross-linked peptides for the ternary protein assembly formed by Bim1, Bik1 and Kar9 identified under condensed (150mM NaCl) **(3A)** and dilute (500mM NaCl) conditions **(3B)**. Crosslinked peptides identified after trypsin proteolysis and filtered for spectrum quality (material and methods) were mapped onto the linear protein sequences with annotated domain and regions. Inter-protein crosslinks (inter-XL) are shown in grey, intra-protein crosslinks (intra-XL, Intralinks) in blue, self-links in orange and inter-XL also detected in partitioning assay in red. The domain abbreviations in the figure are as follows; CH: Calponin Homology domain, EBH: End Binding Homology domain, CAP-Gly: Cytoskeleton-Associated Protein-Glycine-rich domain, Zn-F: Zinc Finger domain, SR: Spectrin Repeat.

Remarkably, comparing the crosslinks identified upon trypsin proteolysis between the proteins in dilute phase and those in the condensed phase, shows that Kar9 P1 and P2 (highlighted in red in Figure 3A) engage with Bik1 exclusively when in the condensed phase (Figure 3A, 3B, S3C, S3D). These interactions were concentrated around the first 30 residues of Bik1 coiled-coil domain. In the dilute phase, P1 engaged in intramolecular interactions instead (two intra-XL 462-562 and 462-564 were identified only in dilute phase), suggesting that sequences of the disordered domain engage these patches and may act as competitors of Bik1. This observation is in line with our FRAP experiments, where the full-length C-terminal domain (fragment 1) recovered more rapidly (measured by the *t_1/2_* value) but less extensively (the time to full recovery, Figure 1C) than fragment 6. The latter probably reflects the fact that fragment 1 seems to have several Bik1-interaction interfaces (Figure 1B). In contrast, the former suggests that at least the interaction interface present in fragment 6, which is also present in fragment 1, interacts with Bik1 more dynamically in the context of the entire C-terminal domain than in isolation. We speculate that elements of this Kar9 domain might compete with Bik1 for interaction and result in this dynamic behavior.

Further analysis of the cross-linking data identified specifically 10 additional cross-links between Kar9 and Bik1 upon phase separation. In contrast, only 2 cross-links were detected between Kar9 and Bik1 in the dilute phase (one in the structured part and one in the C-terminal domain of Kar9). These data further support the notion that additional interactions link Kar9 and Bik1 in condensates. Strikingly however, our cross-linking data suggest that most interactions in the ternary condensate involved Bik1 and Bim1. Consistently fewer inter-protein cross-links were observed in the dilute phase, while intra-protein cross-links were comparatively more abundant (Figure S3A). Importantly, protein abundance remained nearly consistent across conditions (Figure S3B). Thus, whereas Bik1, Kar9, and Bim1 directly interact with each other in the dilute phase, the increase of local protein concentration upon phase separation reveals the existence of many more interactions sites. P1 and P2 in Kar9 are only two of a larger network of interactions between Kar9 and Bik1. Nevertheless, the strong impact of mutating them on partitioning of the entire disordered region indicates that the avidity that they provide is instrumental for Kar9-Bik1 co-condensation (Figure 2C).

Furthermore, the fact that P1- and P2-mediated Kar9-Bik1 interactions are exclusively detected in the condensed phase suggests that they are probably of low-affinity (Figure 3A, 3B, S3C, S3D). Taken together, both the XL-MS and partitioning assay converge to pinpoint that P1 and P2 are important for interaction with Bik1, and further validate our partitioning assay for directly identifying low-affinity interactions and their molecular determinants.

### 4. Kar9 interacts with Bik1 in vivo via P1 and P2

Next, we asked whether P1 and P2 contribute to Kar9-Bik1 interaction in vivo as well. Thus, we mutated the positive charges of these patches to alanines in the yeast genome and tagged the mutated proteins with mNG, generating the alleles *kar9-562aDa-*mNG aka *kar9-p1*^−^, and *kar9-579aDa*-mNG aka *kar9-p2*^−^. We then imaged these mutant cells and measured Kar9 levels at cytoplasmic microtubule plus-ends in pre-anaphase cells (Figure 4A). Mutating either P1 or P2 reduced Kar9 levels at the plus-ends by about 20-30% each (Figure 4A). The effect of these mutations was not additive with that of removing the Bik1 protein altogether (*bik1*Δ mutant background) (Figure 4A). Thus, these data are consistent with P1 and P2 of Kar9 interacting with Bik1 in vivo contributing thereby to Kar9 recruitment to microtubule tips. Consistent with the in vitro data, mutating both patches at the same time (*kar9-562aDa-579aDa*-mNG, i.e., *kar9-p1p2*^−^) decreases further the amount of Kar9 at microtubule tips by about 60% (Figure 2C, 4A). Thus, P1 and P2 act synergistically in Kar9 recruitment to microtubule tips.

**Figure 4:**
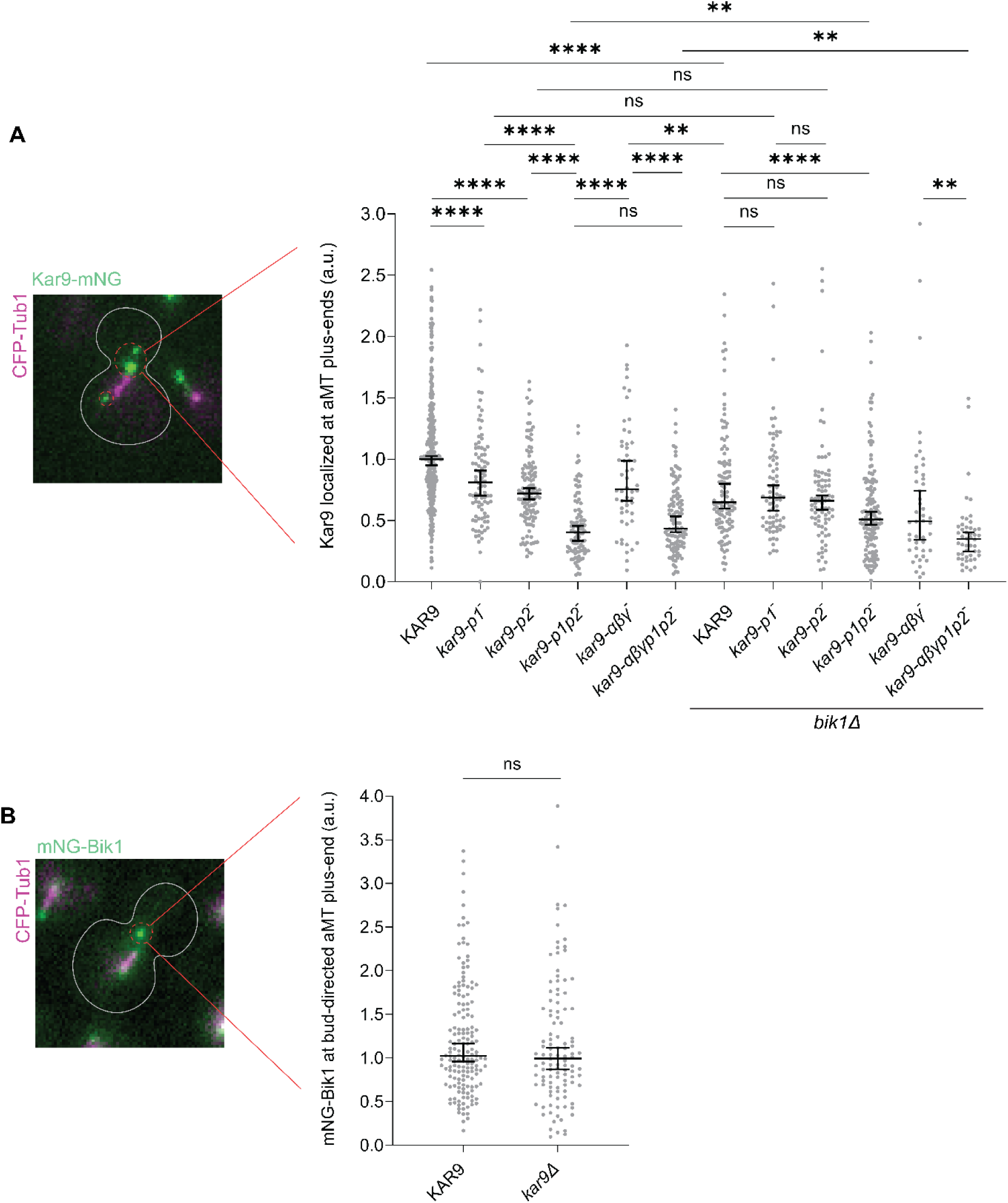
Kar9-Bik1 interaction in vivo. **4A:** Localization of Kar9 variants at cytoplasmic microtubule plus-ends. Each dot represents localization within a single cell; Mann-Whitney test; ns: non-significant, *p<0.05, **p<0.01, ***p<0.001, ****p<0.0001. Bars represent median and standard deviations. **4B:** Localization of mNG-Bik1 at cytoplasmic microtubule plus-ends in presence and absence of Kar9. Each dot represents localization within a single cell; Mann-Whitney test; ns: non-significant. Bars represent median and standard deviations.

Notably, the fluorescence intensity of Kar9-p1p2^−^ to microtubule tips was slightly restored in the *bik1*Δ mutant cells compared to the wild type control (Figure 4A). This suggests that in the absence of P1 and P2, Bik1 rather excludes Kar9 from microtubule tips instead of recruiting it.

Kar9 has been shown to undergo self-oligomerization using α, β, and γ interfaces (Kumar *et al*., 2021; Meier *et al*., 2023). Interestingly, no additive effect was observed upon combining the p1^−^ and p2^−^ mutations with the mutations affecting Kar9 oligomerization (i.e., *kar9-αβγp1p2*^−^; Figure 4A). The levels of Kar9- αβγp1p2^−^ on microtubules remained similar to those of Kar9-p1p2^−^, which are lower than those of Kar9-αβγ^−^. These observations suggest that Kar9 oligomerization mediated by the α, β and γ interfaces takes place subsequent to Kar9 recruitment to the +TIP body, in a P1- and P2-dependent manner. P1 and P2 may be required for the local concentration of Kar9 to pass its critical concentration for oligomerization. Remarkably, whereas the recruitment of Kar9-αβγ^−^ to the +TIP body strongly depends on Bik1, that of Kar9-αβγp1p2^−^ does not, confirming that P1 and P2 mediate the effects of Bik1 in Kar9 recruitment. Importantly, cells lacking Kar9 still localized Bik1 to microtubule tips indistinguishably from wild type (Figure 4B). Thus, whereas Bik1 facilitates Kar9 recruitment to microtubule ends, Kar9 does not contribute to Bik1 recruitment. This underlines that ubiquitous +TIPs mediate the recruitment of patterning +TIPs and not vice-versa.

### 5. Kar9-Bik1 interactions influence properties of the +TIP body and mitotic spindle positioning

Our in vitro and in vivo data identified at least two interfaces mediating Kar9 interaction with Bik1 and showed that these interfaces play an important role in vivo. Based on this, we next wanted to know whether these interactions affect the material properties of +TIP body. Mixing Kar9 into Bik1 droplets decreased their capillary velocity as measured upon droplet fusion in vitro (Figure 5A, Y-axis is inverse capillary velocity), supporting the notion that Kar9 may modulate the material behavior of the +TIP body. Additionally, the Kar9-Bik1 droplets showed biphasic behavior with appearance of condensed clusters within bigger droplets, unlike Bik1 droplets (Figure 5A). Unable to produce Kar9-p1p2^−^ recombinantly, we could not test whether the P1 and P2 patches contribute to Kar9’s effect on Bik1 droplets in vitro. Instead, we asked whether P1 and P2 affected the behavior of the +TIP body in vivo.

**Figure 5:**
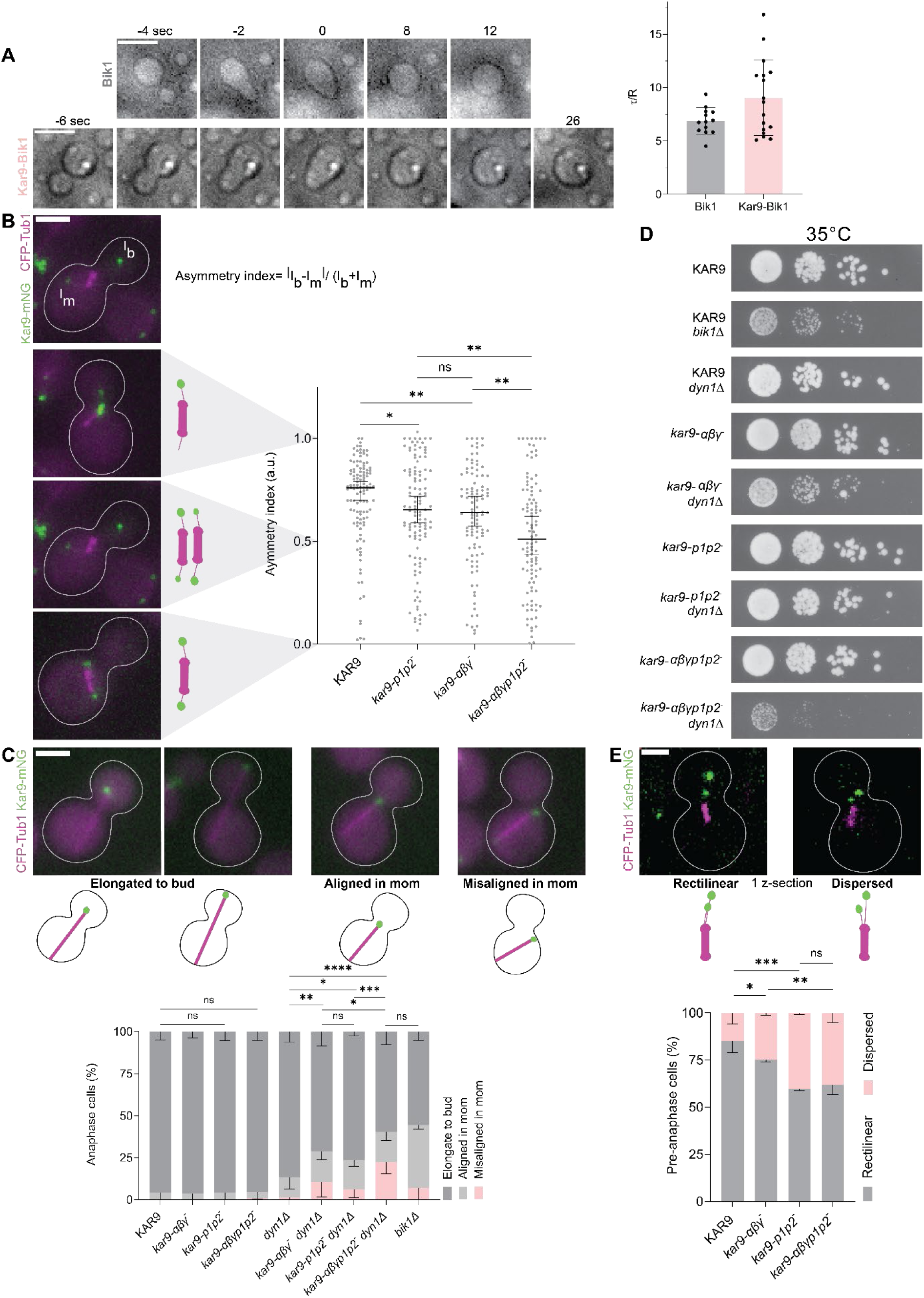
Impact of Kar9-Bik1 interactions on +TIP body’s properties and cell viability. **5A:** Left panel shows representative movies of Bik1 and Kar9-Bik1 droplets relaxing after fusion. Time point 0 indicates the frames with fused droplets having uniaxial deformation. Right panel has τ/R (i.e., inverse capillary velocity) measurements from Bik1 and Kar9-Bik1 droplets with each dot representing one fusion event. Error bars represent standard deviations and histograms indicate mean. Scale bar: 2 µm. **5B:** Quantification of +TIP body asymmetry in pre-anaphase cells, each dot represents quantification from one cell; Mann-Whitney test; ns: non-significant, *p<0.05, **p<0.01, ***p<0.001,****p<0.0001. **5C:** Analysis of alignment of anaphase spindles, error bars represent standard deviations. P values compare ‘Elongated to bud’ and the mispositioned categories (‘Aligned in mom’ and ‘Misaligned in mom’) determined by two proportion z-test; ns: non-significant, *p<0.05, **p<0.01,***p<0.001, ****p<0.0001. Scale bar: 2 µm. **5D:** Spot assay to check viability of strains harboring different Kar9 variants at 35°C. **5E:** Quantification of +TIP body conformation in pre-anaphase cells, error bars represent standard deviations. Rectilinear conformation across strains compared by student’s t-test (2-tailed and unpaired with unequal variance); ns: non-significant, *p<0.05, **p<0.01, ***p<0.001, ****p<0.0001. Scale bar: 2 µm.

In metaphase, the asymmetric localization of Kar9 to only one spindle asters (Liakopoulos *et al*., 2003; Maekawa *et al*., 2003), which is established shortly after spindle assembly (Meziane, Genthial and Vogel, 2021), is facilitated by homotypic Kar9 multivalency (Kumar *et al*., 2021; Meier *et al*., 2023), suggesting that phase separation contributes to symmetry breaking, possibly through Ostwald ripening of the +TIP body (Meier, Steinmetz and Barral, 2024). Therefore, we asked whether Kar9-Bik1 interaction also contributes to symmetry breaking. Remarkably, the *kar9-p1p2*^−^ mutant cells showed an increase in Kar9 symmetry, to a similar level as that observed in the cells expressing a Kar9 variant defective in homotypic oligomerization (*kar9-αβγ*^−^ mutant cells; Figure 5B; Meier *et al*., 2023). To determine whether this defect simply reflects the role of P1 and P2 in recruiting and locally increasing Kar9 concentration, and hence, promoting its oligomerization, we asked whether the effect of the p1p2^−^ mutation is epistatic with that observed upon erasing the α, β and γ interfaces. Strikingly, unlike what we observed for Kar9 recruitment to microtubule ends, the Kar9-αβγp1p2^−^ mutant protein showed a stronger loss of asymmetry compared to the Kar9-αβγ^−^ and Kar9-p1p2^−^ variants (Figure 5B). Thus, the interaction of Kar9 with Bik1 through the interfaces P1 and P2 is required on its own for symmetry breaking, in addition to Kar9 self-interaction. This observation is consistent with P1 and P2 modulating the material properties of the +TIP body.

To explore further whether Kar9-Bik1 interaction indeed affects the material properties of the +TIP body, we asked whether inactivating P1 and P2 had any effect on the capacity of the +TIP body to harvest and transmit the forces generated by microtubule shrinkage at the cell cortex and move the spindle towards the bud during anaphase. Hence, we investigated whether the *kar9-p1p2^−^* mutant cells showed spindle alignment defects in late mitosis. In anaphase, both Kar9 and the microtubule-dependent, minus-end directed motor-protein dynein contribute independently of each other to pulling one of the two poles of the elongating spindle into the bud (Miller and Rose, 1998; Morelli *et al*., 2025). As a consequence, inactivating Kar9 or dynein individually has only minor effects on spindle elongation, whereas inactivating both causes spindles to elongate in the mother cell. These cells generally fail to segregate a daughter nucleus into the bud, form multinucleated mother cells and are unable to generate viable daughters (Meier *et al*., 2023). Therefore, simultaneously inactivating both pathways is lethal. Consistent with these pathways mutually compensating for each other, the dynein defective, *dyn1Δ* single mutant cells elongated their spindle properly during anaphase (Figure 5C), as did the *kar9-p1p2*^−^, *kar9-αβγ ^−^* and *kar9-αβγp1p2^−^* mutant cells. However, a substantial fraction of the *kar9-p1p2*^−^ *dyn1Δ*, *kar9-αβγ ^−^ dyn1Δ* and *kar9-αβγp1p2^−^ dyn1Δ* double mutants cells failed to elongate the spindle along the mother-bud axis and into the bud (Figure 5C). This demonstrates that mutating the homotypic and heterotypic interaction interfaces of Kar9 interfered with the role of the Kar9 +TIP body in pulling the spindle into the bud in the absence of dynein, as already demonstrated for the αβγ interfaces (Meier *et al*., 2023; Morelli *et al*., 2025). Remarkably, here again the effects of abrogating the homotypic and heterotypic interaction interfaces were additive with each other, indicating that both types of interactions contribute to generating the material properties necessary for transmitting the forces generated by microtubule shrinkage at the cell cortex. Accordingly, whereas the *dyn1Δ, kar9-p1p2*^−^, *kar9-αβγ*^−^*, and kar9-αβγp1p2*^−^ single mutants grew well at all physiological temperatures (24-35°C), deleting the dynein gene in the *kar9* mutant strains affected their ability to grow at 35°C (Figure 5D, S5C). The defect was strongest for the *kar9-αβγp1p2*^−^ allele. Thus, the homotypic and heterotypic interaction interfaces are both required for proper Kar9 function, and they function at least partially independently of each other.

Importantly, the *kar9-p1p2*^−^, *kar9-αβγp1p2*^−^ or *bik1Δ* mutant cells showed no defect in mitotic spindle positioning and alignment along the mother-bud axis in the mother cell (Figure S5A). At that stage, force generation does not depend on microtubule depolymerization but on the activity of the motor protein Myo2 (Kumar *et al*., 2021). Thus, even when Kar9 recruitment is reduced by ∼60%, the +TIP body still transduces the Myo2-generated forces properly to the spindle.

Interestingly, cells with *kar9-p1p2*^−^ frequently showed an increased number of +TIP body foci in comparison to the wild type (Figure S5B). Wild type pre-anaphase cells generally contained only one +TIP body (46% of the cells). In the corresponding *kar9-p1p2*^−^ mutant cells, this number dropped to 19%. The *kar9-αβγ*^−^ mutant cells showed a similar phenotype. However, this phenotype was not additive upon combining these mutations. These observations suggest that perturbing Kar9-Kar9 and Kar9-Bik1 interactions affects the ability of +TIP bodies to fuse. Furthermore, in wild type cells harboring several +TIP bodies, these tend to be deposited in a straight line including the SPB, indicating that they are all deposited on the same microtubule or bundle thereof ((Morelli *et al*., 2025; Figure 5E). This was less the case in the *kar9-p1p2*^−^and *kar9-αβγ*^−^ mutant cells. Together, these observations highlight the importance of the multivalent, redundant interactions of Kar9 with the ubiquitous +TIP Bik1 in determining the behavior of the +TIP body, most likely by modulating its material properties.

### 6. Kip2 and Kar9 interact with Bik1 through similar patches

Keeping the interactors of the +TIP body in mind, we noticed Kip2, a microtubule plus-end directed kinesin and microtubule polymerase, also harbors similar patches like P1 and P2 in Kar9. Kip2 accumulates in the Kar9 +TIP body at the plus-end of cytoplasmic microtubules (Chen et al., 2023; Carvalho et al., 2004). Studies have shown that Kip2 accumulation requires it to interact with Bik1 (Carvalho *et al*., 2004; Chen *et al*., 2023). Furthermore, interaction of Kip2 with Bik1 was mapped to the 60 amino acids long, C-terminal disordered tail of Kip2 (Kip2 tail/T) (Chen *et al*., 2023). Truncated versions of Kip2 from which its motor domain (Kip2-NMD) is removed still localized to the plus-end of microtubules in a Bik1-dependent manner and this localization is lost upon removing the Kip2 tail. Interestingly, we noticed that the Kip2 tail comprised patches of oppositely charged amino acids, 670-KDK and 684-RDR (hereon, referred to as Q1 and Q2) (Figure 6A). Like Kar9 where P1 and P2 are separated by 15 amino acids, Q1 and Q2 in Kip2 are separated by 13 amino acids. Upon manually aligning P1, P2, Q1 and Q2 across fungal species, we found presence of conserved hydrophobic residues (L/I/V) surrounding K/R-D/E-K/R patches, although their exact position is not consistent between P1/P2 and Q1/Q2 (Figure S6A). Furthermore, we noted presence of a serine in −6 of the first K/R in all cases except Q1. These suggested that Kip2 and Kar9 may interact with Bik1 through similar mechanisms. Although the surrounding hydrophobic residues don’t share positional conservation between the two proteins, they might still assist in the interaction with the Bik1 coiled-coil given their presence in the disordered regions.

**Figure 6:**
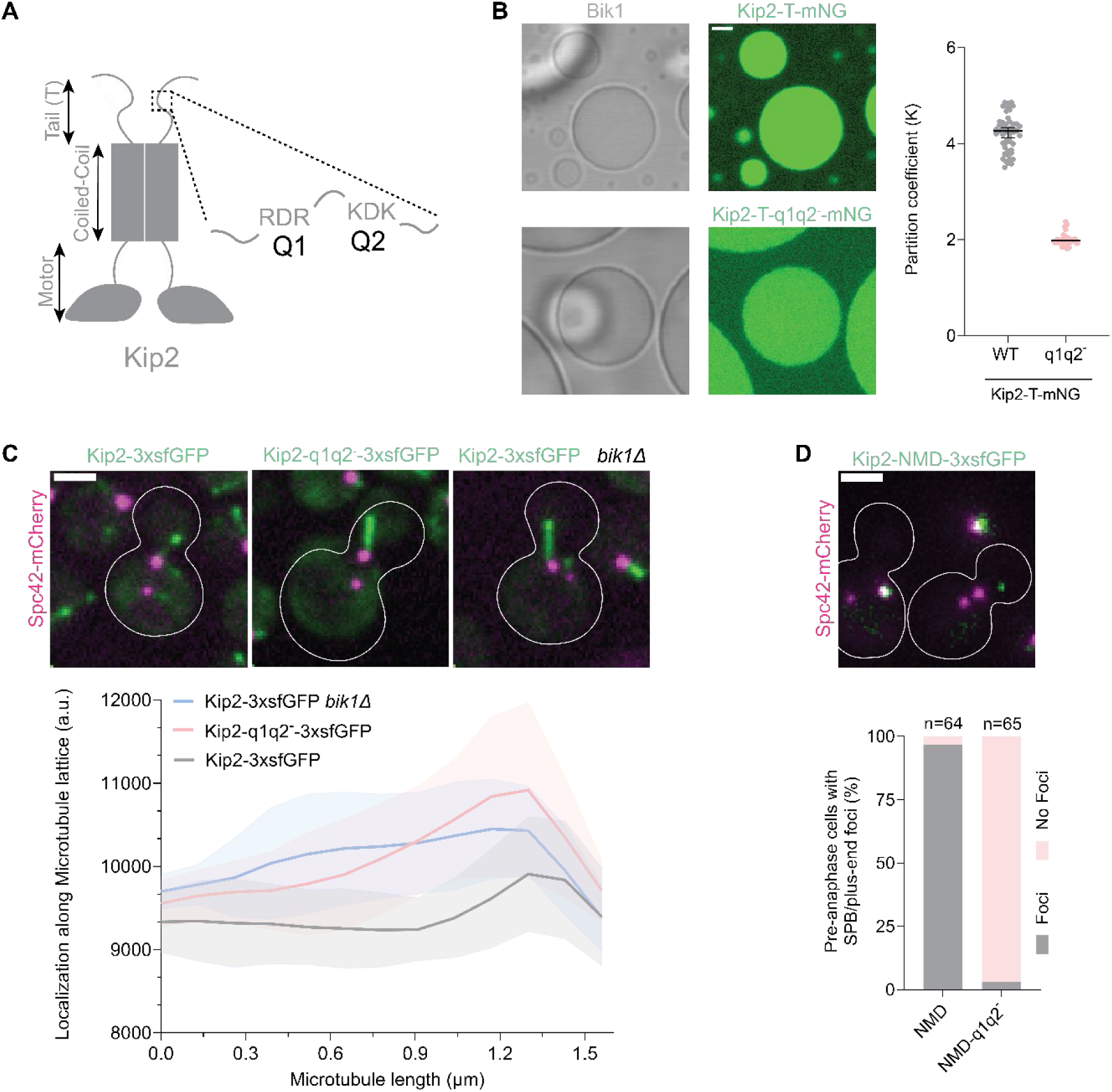
Bik1’s interaction with Kip2 follows similar grammar. **6A:** Representation of Kip2 with alternating charged motifs, Q1 and Q2. **6B:** Partitioning behavior represented by partition coefficients of Kip2 tail variants into Bik1 droplets. Each dot represents K in a single droplet. Scale bar: 2 µm. Presence of full-length fragments were verified by Western Blot (Figure S6D). **6C:** Localization of Kip2-3xsfGFP in presence and absence of Bik1, and Kip2-q1q2^−^-3xsfGFP along the microtubule lattice in pre-anaphase cells. Each bold line represents mean profile of at least 7 microtubules, with standard deviations represented as shaded area. Represented images are in sum projection Scale bar: 2 µm. **6D:** Percentage of pre-anaphase cells with a +TIP body focus of Kip2-NMD-3xsfGFP. Represented image is in max projection. Scale bar: 2 µm.

Supporting this idea, the tail of Kip2 C-terminally tagged with mNG (Kip2-T-mNG) partitioned into Bik1 droplets above the levels reached by the mNG control (Figure 6B, 1B). The partition coefficient K∼4 of Kip2 tail suggests low-affinity interaction with Bik1, similar to Kar9.

Mutating the positively charged residues of Q1 and Q2 to alanines (Kip2-T-670-aDa-684-aDa-mNG, aka Kip2-T-q1q2^−^-mNG), abolished this enrichment. Thus, Q1 and Q2 drive Kip2-T interaction with Bik1 in vitro. In vivo studies indicated that this was the case in cells as well.

Indeed, the Kip2-670aDa-684aDa (aka Kip2-q1q2^−^) tagged with three copies of super-folder GFP (sfGFP), Kip2-q1q2^−^-3xsfGFP, accumulated less prominently at microtubule plus-ends than the tagged wild type protein and rather spreads along the lattice of cytoplasmic MTs, like reported for Kip2-ΔT-3xsfGFP and as we see for Kip2-3xsfGFP in *bik1Δ* mutant cells (Figure 6C). Likewise, unlike Kip2-NMD-3xsfGFP, Kip2-NMD-q1q2^−^-3xsfGFP failed to decorate microtubule tips (Figure 6D). Even in the cells (approximately 40%) that showed Kip2-NMD-q1q2^−^-3xsfGFP clusters, these did not localize to the +TIP body (Figure S6B).

Taken together, these observations confirm that Q1 and Q2 in Kip2 utilize similar mechanisms as P1 and P2 in Kar9 to interact with Bik1. Furthermore, our data highlight the role of the ubiquitous +TIP Bik1 in modulating the molecular environment of microtubule plus-end by stabilizing their interactors with specialized contributions to microtubule dynamics and function.

## Discussion

Although composed of tubulin dimers drawn from the cytoplasmic pool, different microtubules execute different functions in the same cell, depending on where they are nucleated (Chen *et al*., 2019) or the region of the cell they extend into. For example, mitotic cells comprise kinetochore microtubules, which pull chromosomes towards the spindle poles, interpolar microtubules, which push the two spindle poles away from each other, and cytoplasmic microtubules that move and position the spindle within the cell and instruct the cell cortex about where to assemble the cleavage apparatus. However, the molecular mechanisms cells utilize to specify these distinct behaviors and functions are not well understood. There is accumulation of evidence that the structures decorating microtubule plus-ends, such as +TIP bodies, contribute to specifying microtubule behavior and function (Song et al., 2023; Maan *et al*., 2023; Meier *et al*., 2023; Miesch *et al*., 2023; Morelli *et al*., 2025). Two proteins are at the core of these bodies on most, if not all, microtubules, namely EBs and CLIP-170. Our results indicate that specialized +TIP components, i.e., patterning +TIPs such as Kar9 (APC/SLAIN/MACF), can modulate their behavior to specific context and function. Indeed, at least in the case of Kar9, our data make the following points.

First, our data show that Kar9 interacts with Bik1 through at least three interfaces, lending extensive multivalency between +TIP body components. Previous work (Stangier *et al*., 2018; Czub *et al*., 2025) and our crosslinking mass spectrometry data presented here demonstrate that the ubiquitous +TIPs EB1/Bim1 and CLIP-170/Bik1 are linked through a dense network of homo- and heterotypic interactions, promoting their phase separation on most if not all microtubule plus-ends, from yeast to mammals (Song et al., 2023; Maan *et al*., 2023; Meier *et al*., 2023; Miesch *et al*., 2023; Morelli *et al*., 2025). Already published data establish that Kar9 interacts with itself through at least three independent interfaces and with Bim1 through at least three interfaces as well (Manatschal *et al*., 2016; Meier *et al*., 2023; Kumar *et al*., 2017, 2021). Our study provides additional evidence for multivalent interactions of Kar9 with each core ubiquitous +TIP components and hence supports the idea that Kar9 plays a central role in the organization of the yeast cytoplasmic +TIP body. Importantly, erasing the interfaces between Kar9 and Bim1 (Manatschal *et al*., 2016) and between Kar9 and Bik1 (this study) have strong effects on the recruitment of Kar9 to the +TIP body and on +TIP body function. These observations strengthen the notion that Kar9 recruitment and the assembly of the +TIP body are a cooperative process, consistent with being driven by phase separation. In other words, all these interfaces contribute to setting the critical concentration for phase separation to a low level.

Second, our study narrows down two interfaces in Kar9 and identifies thereby a K/R-D/E-K/R-centered short linear motif (SLiM) driving the interaction of Kar9 with Bik1. Remarkably, this SLiM motif is repeated in Kar9, conserved across Kar9 homologs, even if not always placed at the same position, and utilized by the kinesin Kip2 as well. We would like to make three points about this SLiM. i) The conserved nature of this motif suggests that it interacts with some folded part of Bik1, as supported by the crosslinking experiments, which suggest interaction of Kar9 SLiM motifs with the N-terminal extremity of the long coiled-coil domain of Bik1. This position complexifies studying the Bik1 side of this interaction, as mutations in the coiled-coil is likely to affect Bik1 dimerization as well. Structural analysis will be needed to pinpoint mutations that affect Bik1 interaction with Kar9 and Kip2 without interfering Bik1 dimerization. ii) This motif involves a combination of charged and potentially hydrophobic and serine/threonine residues, suggesting that it does not involve solely electrostatic mechanisms of binding. Future structural studies will also clarify how the motif found in Kar9 and Kip2 engages in interactions with Bik1. It will be also interesting to determine whether this motif is found in other interactors of Bik1. Remarkably, we were so-far unable to identify any convincing occurrence of this SLiM in Cik1 and Kar3, two key interactors of Bim1 and Bik1 in the yeast spindle (Molodtsov *et al*., 2016; Kornakov, Mollers and Westermann, 2020; Kornakov and Westermann, 2023). Thus, an attractive possibility would be that the SLiM is only found in Kar9 and Kip2 and thus is specific to the +TIP body of cytoplasmic microtubules responsible for mitotic spindle positioning. For example, if some regulatory modification of Bik1 inhibits or promotes the exposition of the relevant interface in specific subcellular contexts, this could contribute to specifying the composition and behavior of the affected +TIP bodies. In that respect, it is interesting that inactivating two SLiMs in Kar9 caused its displacement from the cytoplasmic +TIP body, in a Bik1-dependent manner. Hence, Bik1 seems to serve as much as a platform for the recruitment of interactors as a shield that restricts access to non-interacting proteins. This may be an important mechanism for diversifying +TIP bodies, possibly explaining for example why Bik1 recruits Kar9 on cytoplasmic but not on spindle microtubules. iii) The SLiM identified in this study may be subject to cis-regulation whereby it is inhibited to interact with Bik1, given one of the two copies of the SLiM interact with another region in the disordered C-terminal domain of Kar9, as suggested by our crosslinking experiments. Possibly, this intramolecular Kar9-Kar9 interaction competes with Bik1. Supporting this idea, the full-length C-terminal domain of Kar9 interacts more dynamically with Bik1 than the isolated SLiM containing Kar9 fragments, despite the enhanced avidity provided by the presence of several copies of the SLiM in the former. We propose that this dynamic behavior is enhanced by such competitions. iv) Whereas the SLiM repeats of Kar9 affect the properties of the +TIP body, at first approximation those of Kip2 may simply promote the retention of this microtubule polymerase to microtubule tips.

Third, our in vitro and in vivo data indicate that the Kar9-Bik1 interaction contributes to tuning the material property of the +TIP body to promote proper function. In vivo, this manifests in at least two ways: the asymmetry of Kar9 distribution on the pre-anaphase spindle and the movement of the spindle into the bud upon metaphase onset. Indeed, previous work showed that any perturbation affecting Kar9 dimerization (Kumar *et al*., 2021) and Kar9 oligomerization (Meier *et al*., 2023) and hence, conditions that affect the ability of Kar9, Bik1, and Bim1 to phase separate in vitro, delay or prevent symmetry breaking. Likewise, preventing pre-anaphase phosphorylation of Kar9, a process regulating phase separation of Kar9, delays symmetry breaking (Liakopoulos *et al*., 2003; Hotz *et al*., 2012; Kumar *et al*., 2021). Together, these data indicate that symmetry breaking is driven by phase separation, possibly through Ostwald ripening (Meier, Steinmetz and Barral, 2024). Our data suggest that the Kar9-Bik1 interaction promotes this process. It might do so by its role in promoting Kar9 recruitment and facilitating Kar9-Kar9 oligomerization. Since mutating P1 and P2 as well as the three Kar9-Kar9 homotypic interfaces pinpoints their requirement for a robust asymmetry of Kar9 on the spindle, we propose that both Kar9 recruitment by Bik1 and its subsequent oligomerization contribute to symmetry breaking. Furthermore, upon anaphase onset, Kar9 starts to promote the movement of the spindle into the bud. Previous studies have demonstrated that this process involves harvesting the forces generated by microtubule shrinkage at the bud cortex (Morelli *et al*., 2025). In order to couple the force dissipated by microtubule shrinkage, the Kar9 +TIP body needs to maintain interactions with the cell cortex as well as with shrinking microtubules. Here again, Kar9 and its ability to self-interact are crucial. Our data now indicate that Kar9 interaction with Bik1 is also instrumental. Based on our current in vitro data, we propose that the decrease in capillary velocity conferred by Kar9 to Bik1 droplets may be pivotal in the process of force transduction during microtubule shrinkage. However, future work will be required to test this idea and characterize more precisely the mechanism of force coupling within the +TIP body. Finally, the observation that affecting Kar9 interaction with Bik1 may interfere with microtubule bundling in the cytoplasm is very intriguing as well. This observation might reveal a role of Kar9 in promoting the fusion of +TIP bodies upon their contact, which will be interesting to test in the future.

On a more general level, our data reveal the importance of identifying and dissecting the highly dynamic, low-affinity interactions taking place between condensate components. While these dissections can be tedious and generally remain incomplete, they enable one to characterize the mechanisms driving condensate formation, modulating the material behavior of the condensate, and deciphering how these features contribute to the biological function of condensate components and the condensate itself in living cells. Low-affinity interactions are notoriously difficult to identify and measure, even before dissecting their underlying molecular determinants. The partitioning-based assay that we introduced here proved very effective in its simplicity for this purpose. The generalization potential that it offers suggests that it might be a solution of choice for identifying and dissecting the molecular determinants of phase separating processes.

## Materials and Methods

### Protein overexpression

Kar9 variants tagged with mNeonGreen in the C-terminus and 6xHis-tag in the N-terminus were cloned into pACEMBLE-A vector by homologous recombination in *E. coli* DH5α. Recombinant plasmid was transformed into *E. coli* BL21 (RIL). Transformed *E. coli* was grown overnight (o/n) at 37°C until saturation, followed by inoculation into fresh media with 1% V/V inoculum. Once culture reached log phase (OD 0.4-0.8), 1mM IPTG was added to the culture and incubated for o/n at 20°C. Cells were harvested by spinning at 4000 rpm for 15 minutes.

### Protein purification and preparation

The DNA encoding the full-length *S. cerevisiae* Bik1 (Uniprot ID: P11709) was cloned into the pET-based bacterial expression vector PSPCm2, which encodes for an N-terminal 6×His-tag followed by a PreScission cleavage site, using a positive selection method (Olieric *et al*., 2010). The Bik1 clone has been described in our previous study (Meier *et al*., 2023).

Plasmids verified by sequencing were transformed into *E. coli* strain BL21-CodonPlus (DE3)-RIPL (Agilent #230280). For protein production, culture was grown in LB medium containing 50 μg/ml of kanamycin at 37 °C until reaching an OD_600_ of 0.6, then cooled down to 18 °C. Subsequently expression was induced with 0.75 mM isopropyl 1-thio-β-D-galactopyranoside (IPTG), and the induced cells were incubated in a shaker for 16 h at 18 °C. Following harvesting by centrifugation, the cells were lysed by sonication in the presence of protease inhibitors (cOmplete cocktail; Roche), 0.1% bovine deoxyribonuclease I and 1 mM phenylmethylsulfonyl fluoride (PMSF) in the lysis buffer (20 mM Tris-HCl, pH 7.5, supplemented with 500 mM NaCl, 10 mM imidazole and 2 mM β-mercaptoethanol).

Proteins were purified by immobilized metal-affinity chromatography (IMAC) on a HisTrap HP nickel-Sepharose column (GE Healthcare) at 4 °C following the manufacturer’s instructions. The column was equilibrated in IMAC buffer A (20 mM Tris-HCl, pH 7.5, supplemented with 500 mM NaCl, 10 mM imidazole, and 2 mM β-mercaptoethanol). Proteins were eluted by IMAC buffer B (IMAC buffer A containing 400 mM imidazole in total).

Protein samples were further loaded onto a size-exclusion chromatography (SEC) HiLoad Superdex 200 16/60 column (GE Healthcare) equilibrated in SEC buffer (20 mM Tris-HCl, pH 7.5, supplemented with 500 mM NaCl, 10% glycerol, and 1 mM DTT) or in XL buffer (10 mM HEPES-NaOH, pH 7.5, supplemented with 500 mM NaCl and 1 mM DTT) for XL-MS experiments. Bik1 protein-containing fractions were pooled and concentrated to the desired concentration. The protein quality and identity were assessed by SDS-PAGE and mass spectrometry, respectively.

### Partitioning assay

50 OD (1 ml of culture with OD 1.0 at 600 nm would result to 1 OD) cells were resuspended with 1 ml cold PBS and spun at 14000 rpm for 1 min at 4°C. Supernatant was removed and cell pellet was lysed with 500 µl lysis buffer (50 mM Tris, 500 mM NaCl, 0.5 mM EDTA, 1 mM MgCl_2_, 0.2% Triton X-100, 10% glycerol, pH 7.5) freshly supplemented with 2X protease inhibitor cocktail (cOmplete^™^, EDTA-free Protease Inhibitor Cocktail, Catalogue no.-11873580001) and 400 µl (approximately measured in Eppendorf) glass beads (Accessories Beads Glass beads, 0.5 mm, Carl Roth) in a FastPrep^®^-24 5G bead beating grinder and lysis system bead beater at 6 m/sec with QuickPrep adapter for 4 cycles at 4°C as following: 50 sec beating followed by 2 min pause followed by 50 sec beating followed by 1 min pause followed by 50 sec beating followed by 2 min pause followed by final 50 sec beating. Lysed cells were centrifuged at 15000 rpm, 4°C for 15 minutes. Lysates were separated, flash frozen in liquid N_2_ and stored at −80°C. Size of recombinant products were verified by western blot with N-terminal binding Anti-His antibody (Figure S1, S2A, S6C). Lysate accounting to 27% of the reaction volume was premixed with 30 µM purified recombinant Bik1, and phase separation was induced with equal volume of no salt containing buffer (20 mM Tris, 1mM DTT, pH 7.5) to reach final salt concentration of 250 mM NaCl. Droplets were induced on chambered coverglass (Nunc™ Lab-Tek™ II Chambered Coverglass, catalogue number 155409) and imaged approximately 4 min post induction in confocal settings with Zeiss Airyscan microscopy using 2% 488 nm laser power with gain of 380, T-PMT settings and stacks (stack size-0.38 µm). Imaging was initiated from the bottom of the slide and 7^th^ stack was used for analysis of partition coefficients. Mean intensity within a selected region of interest in the droplet and its surrounding was quantified. Ratio of intensity inside (I_in_) to outside (I_out_) resulted in partition coefficient.

### Fluorescence Recovery After Photobleaching

Samples were prepared as in the partitioning assay. Fluorescence of the sample was recorded after 5 min 30 s post induction in a single confocal plane with 488 nm laser at 20% power using a Nikon Spinning Disk SoRa system for 2 s. 2 µm*2 µm feret of interests were bleached with a FRAP488 laser pulse at 70% power for 150 ms. Fluorescence recovery was monitored for 2 min without delay. For analysis, mean fluorescence intensities in a region of interest within the photo-bleached region of droplets and an unbleached nearby reference droplet were measured for the whole course of imaging. To account for photobleaching during imaging, intensity of the reference droplet was used to normalize the intensity of the photobleached region. Normalized intensities were plotted with time. The data were fit using one-phase exponential association followed by a plateau model in GraphPad Prism 10 and half-lives (t_1/2_) were extracted from the fit.

### Crosslinking reaction and proteomics sample preparation

An equimolar mixture of purified Bim1, Bik1, and Kar9 (∼1 mg/mL) was incubated in 10 mM HEPES buffer (pH 7.5) supplemented with either 150 mM or 500 mM NaCl to generate dilute or phase-separated conditions, respectively. All crosslinking experiments were performed in triplicate, following previously established protocols [PMID: 24356771, PMID: 39885130].

Samples were crosslinked with 1 mM isotopically labeled disuccinimidyl suberate (DSS-d0, DSS-d12; Creative Molecules Inc.) for 30 minutes at 37 °C, then quenched with 0.1 M ammonium bicarbonate for 30 minutes at room temperature. Following quenching, samples were dried in a SpeedVac, resuspended in 8 M urea, reduced with 5 mM Tris(2-carboxyethyl) phosphine (TCEP) for 30 minutes at 37 °C, and alkylated with 10 mM iodoacetamide for 30 minutes in the dark at room temperature. Proteolytic digestion was performed with trypsin or GluC sequencing grade (Promega) at a 1:50 E:S and quenched after ∼16 hours with 5% formic acid. Resulting peptides were purified using C18 columns (The Nest Group) according to the manufacturer’s protocol. Crosslinked peptides were enriched by size-exclusion chromatography (SEC) using an ÄKTA micro chromatography system (GE Healthcare) equipped with a Superdex 200 Increase 3.2/30 column (Cytiva). SEC fractions were then dried and resuspended in 5% acetonitrile with 0.1% formic acid for mass spectrometry analysis.

### Mass Spectrometry sample acquisition

LC–MS/MS analysis was performed on an Orbitrap QExactive+ mass spectrometer coupled to an Easy-nLC 1000 system (Thermo Fischer Scientific). Peptides were separated using a reverse phase column (75 µm ID × 400 mm New Objective, in-house packed with ReproSil Gold 120 C18, 1.9 µm, Dr. Maisch GmbH) across a 60 min gradient: from 5% to 25% in 55 min and from 25% to 40% in 5 min (buffer A: 0.1% (v/v) formic acid; buffer B: 0.1% (v/v) formic acid, 95% (v/v) acetonitrile). The acquisition was performed with one MS1 scan followed by a maximum of 20 scans for the top 20 most intense peptides (TOP20) with MS1 scans (R = 70,000 at 400 m/z, maxIT = 64 ms AGC=1e5), HCD fragmentation (NCE = 28%), isolation windows (1.5 m/z) and MS2 scans (R = 35,000 at 400 m/z, maxIT = 110 ms, AGC = 5e4ms). A dynamic exclusion of 30 s was applied, and charge states lower than three and higher than six were rejected for the isolation.

### Mass Spectrometry data analysis

Data converted to mzXML format were searched with XQuest/xProphet [Walzthoeni] against a database containing the FASTA sequence of Kar9, Bim1 and Bik1 and decoy sequence for false discovery rate estimation. Crosslinked peptides were included in the analysis if they met the following criteria: a) minimum peptide length of 5 amino acids, b) xQuest ld score >20, c) FDR < 0.2 d) top-ranked hit per spectrum, e) identified by at least two spectra (only intra-XL and intralink peptides). The results were visualized using the xiVIEW web-based tool [PMID: 39237202]. Protein intensities were determined by MaxQuant analysis v2.4.13.0 using default parameters and the FASTA sequence of Kar9, Bim1 and Bik1 [PMID: 22772729].

The entire dataset, including raw data, generated tables, and scripts used for the data analysis are available in the PRIDE repository [PMID: 34723319] with the identifier PXD078814.

### Yeast strains and cloning

*S. cerevisiae* strains were constructed in the S288C background. Yeast strains were created by transformation of DNA fragments amplified by PCR which later integrated into the genome by homologous recombination and/or by crossing, sporulation and tetrad dissection. For the creation of Kar9 mutations, mutant primers were used to amplify KAR9 by PCR as multiple fragments overlapping at the sites of the point mutations. The mutant fragments were transformed into *kar9Δ* strain which later recombined with the genome using homologous sequences to produce the mutants. Same approach was utilized for constructing strains with Kip2 variants. Transformants were verified by PCR, sequencing and microscopy, and spores from tetrad dissection were verified by transfer onto relevant selection plates.

### Live-cell fluorescence microscopy

For imaging Kar9-mNG, cells were grown o/n in a rotating wheel in minimal medium lacking Tryptophan. Cells were harvested at 600 g for 30 s. For strains with CFP-Tub1 and Kar9-mNG, cells were imaged in 17 stacks of 0.25 µm each in a Delta Vision microscope (Cytiva) at 100X magnification with 2*2 binning with 300 msec exposure for CFP and GFP channel each and 100% transmission. Time lapse movies were recorded for 12 time points, with 13 sec intervals between each time point. For imaging Kip2-3xsfGFP and Spc72-mCherry, images were acquired at 60X magnification in visitron spinning disk in stream mode, 100% laser power for both channels, and 50 msec exposure for green channel and 80 msec for red channel. Images were acquired in 17 stacks separated by 0.2 µm. For imaging Kip2-NMD-3xsfGFP, cells were imaged at 100X magnification in 17 stacks of 0.25 µm each in a Delta Vision microscope with 2*2 binning and 300 msec exposure for mCherry and GFP channel each and 100% transmission.

### In vivo protein localization analysis

Images acquired at time point 2 were sum-projected and analyzed for localization of Kar9 at cytoplasmic microtubule plus-ends in metaphase cells (cells with 1-2 µm spindle size). Integrated density of a region of interest was extracted in Fiji encompassing the +TIP body (I_droplet_) or the surrounding cytoplasm (I_background_). I_background_ was subtracted from I_droplet_ and subtracted values at both sides of the spindle were added up to represent the localization of Kar9. All the data were normalized to the median of the wild type. For analyzing localization of Kip2 along the microtubule lattice, metaphase cells were selected and localization profiles from SPBs to the microtubule plus-end along a 5-pixel wide line were extracted from sum-projected images. For analysis of Kip2-NMD-3xsfGFP localization, time point 2 was selected to score the localization at the +TIP body in sum-projected images.

### In vitro droplet fusion analysis

Bik1 droplets are induced at 15 µM Bik1 with 250 mM NaCl in buffer containing 20 mM Tris, 10% Glycerol, pH 7.8. Kar9+Bik1 droplets were induced at 15 µM Bik1, 5 µM Kar9 and 250 mM NaCl in buffer containing 20 mM Tris, 10% Glycerol, pH 7.8. Droplets were allowed to settle for 5 minutes post their induction and movies were acquired for 15 minutes in a Delta Vision wide-field microscope in a single z-plane close to the bottom of the coverslip with 150 msec exposure and time interval of 2 sec between time points. To measure the fusion time τ, we started with uni-axially deformed droplets and measured the major axis (L) and the minor axis (W), as described in Ijavi *et al*., 2021. The parameter A which is (L-W)/(L+W) was calculated over time until the fused droplet relaxes to a sphere by eye. Fitting an exponential curve (y=a*exp (-x/τ) +b) to the time-dependent parameter *A* using MATLAB gave the fusion time τ. The radius R of the relaxed droplet is measured as (L+W)/4 and τ/R values (inverse capillary velocity) are extracted for each fusion event.

### In-vivo phenotypic analysis

For scoring the number of +TIP foci, microtubule bundling and spindle alignment during metaphase and anaphase, timepoint 2 was used. To analyze microtubule bundling, +TIP bodies during metaphase were scanned across all the z-sections and scored for alignment with the spindle. If multiple +TIP bodies could be connected along a single line with the spindle pole, particular cells were counted to have “Rectilinear conformation”; otherwise, they were put in the class of “Dispersed conformation”. Cells with one +TIP body were eliminated from this analysis. For analyzing number of +TIP foci, metaphase cells were similarly scanned through all the z-sections, and the maximum number of observed foci were considered. To score for spindle alignment, sum projected images of metaphase and anaphase cells were used and data were classified into groups.

### Yeast viability assay

Starting from OD_600_ 0.05, haploid yeast strains of the given genotypes were spotted in 10× serial dilutions, by spotting 4 µl from the individual dilutions on YPD plates. They were incubated at the indicated temperatures for 2–3 days.

## Supplementary table 1 (in attachment)

Results from crosslinking analysis of Bim1, Bik1 and Kar9 in dilution and phase separation conditions.

## Supplementary table 2 (in attachment)

List of yeast and bacterial strains used in this study.

## Resource Availability

Requests for additional information and resources should be directed to the lead contact, Yves Barral. All yeast strains generated in this study are available upon request.

## Author Contributions

Madhurima Choudhury: Conceptualization, Data curation, Formal analysis, Validation, Investigation, Visualization, Writing—original draft, Writing—review and editing; Federico Uliana: Data curation, Formal analysis, Validation, Investigation, Writing—original draft, Writing—review and editing; Tarik Grubić: Data curation, Writing—original draft, Writing—review and editing; Mateusz P. Czub: Data curation, Investigation, Writing—review and editing; Ana-Maria Farcas: Data curation, Formal analysis, Investigation, Writing—review and editing; Michel O. Steinmetz: Funding acquisition, Investigation, Resources, Writing—review and editing; Yves Barral: Conceptualization, Funding acquisition, Supervision, Investigation, Resources, Writing—original draft, Writing—review and editing.

## Acknowledgement

We sincerely thank Dr. Alexander Leitner (ETH Zürich) for sharing his expertise in crosslinking strategies and for providing access to instruments and data analysis platforms. We thank Sandro M. Meier for sharing purified Bik1 stock, plasmids, and protocols, and for helpful discussions. We also thank all the Barral lab members for constructive discussions, and ScopeM, ETH Zurich for their help in setting up microscopy experiments.

## References

Akhmanova, A. and Steinmetz, M.O. (2008) “Tracking the ends: A dynamic protein network controls the fate of microtubule tips,” Nature Reviews Molecular Cell Biology, pp. 309–322. Available at: 10.1038/nrm2369.

Akhmanova, A. and Steinmetz, M.O. (2015) “Control of microtubule organization and dynamics: Two ends in the limelight,” Nature Reviews Molecular Cell Biology. Nature Publishing Group, pp. 711–726. Available at: 10.1038/nrm4084.

Banani, S.F. et al. (2017) “Biomolecular condensates: Organizers of cellular biochemistry,” Nature Reviews Molecular Cell Biology. Nature Publishing Group, pp. 285–298. Available at: 10.1038/nrm.2017.7.

Beach, D.L. et al. (2000) The role of the proteins Kar9 and Myo2 in orienting the mitotic spindle of budding yeast.

Beutel, O. et al. (2019) “Phase Separation of Zonula Occludens Proteins Drives Formation of Tight Junctions,” Cell, 179(4), pp. 923–936.e11. Available at: 10.1016/j.cell.2019.10.011.

Carvalho, P. et al. (2004) Cell Cycle Control of Kinesin-Mediated Transport of Bik1 (CLIP-170) Regulates Microtubule Stability and Dynein Activation cytoplasmic linker protein CLIP-170 (Perez et al., 1999) and then observed for some other microtubule-associated proteins (MAPs) (Carvalho et al Many of the TIPs that directly bind MTs also have important roles in controlling plus end dynam, Developmental Cell. Howard and Hyman. Available at: http://www.developmentalcell.

Chen, X. et al. (2019) “Remote control of microtubule plus-end dynamics and function from the minus-end,” eLife, 8. Available at: 10.7554/eLife.48627.

Chen, X. et al. (2023) “The motor domain of the kinesin Kip2 promotes microtubule polymerization at microtubule tips,” Journal of Cell Biology, 222(7). Available at: 10.1083/jcb.202110126.

Czub, M.P. et al. (2025) “Phase separation of a microtubule plus-end tracking protein into a fluid fractal network,” Nature Communications, 16(1). Available at: 10.1038/s41467-025-56468-8.

Dominguez, R. and Holmes, K.C. (2011) “Actin structure and function,” Annual Review of Biophysics, 40(1), pp. 169–186. Available at: 10.1146/annurev-biophys-042910-155359.

Goodson, H. V. and Jonasson, E.M. (2018) “Microtubules and microtubule-associated proteins,” Cold Spring Harbor Perspectives in Biology, 10(6). Available at: 10.1101/cshperspect.a022608.

Hotz, M. et al. (2012) “Spindle pole bodies exploit the mitotic exit network in metaphase to drive their age-dependent segregation,” Cell, 148(5), pp. 958–972. Available at: 10.1016/j.cell.2012.01.041.

Hwang, E. et al. (2003) “Spindle orientation in Saccharomyces cerevisiae depends on the transport of microtubule ends along polarized actin cables,” Journal of Cell Biology, 161(3), pp. 483–488. Available at: 10.1083/jcb.200302030.

Ijavi, M. et al. (2021) “Surface tensiometry of phase separated protein and polymer droplets by the sessile drop method,” Soft Matter, 17(6), pp. 1655–1662. Available at: 10.1039/d0sm01319f.

Kornakov, N., Mollers, B. and Westermann, S. (2020) “The EB1-Kinesin-14 complex is required for efficient metaphase spindle assembly and kinetochore bi-orientation,” Journal of Cell Biology, 219(12). Available at: 10.1083/JCB.202003072.

Kornakov, N. and Westermann, S. (2023) “Systematic analysis of microtubule plus-end networks defines EB-cargo complexes critical for mitosis in budding yeast,” Molecular Biology of the Cell, 34(5). Available at: 10.1091/mbc.E23-02-0054.

Kumar, A. et al. (2017) “Short Linear Sequence Motif LxxPTPh Targets Diverse Proteins to Growing Microtubule Ends,” Structure, 25(6), pp. 924–932.e4. Available at: 10.1016/j.str.2017.04.010.

Kumar, A. et al. (2021) “Structure and regulation of the microtubule plus-end tracking protein Kar9,” Structure, 29(11), pp. 1266–1278.e4. Available at: 10.1016/j.str.2021.06.012.

Liakopoulos, D. et al. (2003) Asymmetric Loading of Kar9 onto Spindle Poles and Microtubules Ensures Proper Spindle Alignment actions with the cortical dynein/dynactin complex (Du-jardin and Vallee, 2002). EB1 acts in concert with the adenomatous polyposis coli (APC) tumor suppressor We show that asymmetric localization of Kar9 to the, Cell. Available at: http://www.cell.com/cgi/content/full/.

Maan, R. et al. (2023) “Multivalent interactions facilitate motor-dependent protein accumulation at growing microtubule plus-ends,” Nature Cell Biology, 25(1), pp. 68–78. Available at: 10.1038/s41556-022-01037-0.

Maekawa, H. et al. (2003) “Yeast Cdk1 translocates to the plus end of cytoplasmic microtubules to regulate bud cortex interactions,” The EMBO Journal, 22(3), pp. 438–449. Available at: 10.1093/emboj/cdg063.

Manatschal, C. et al. (2016) “Molecular basis of Kar9-Bim1 complex function during mating and spindle positioning,” Molecular Biology of the Cell, 27(23), pp. 3729–3745. Available at: 10.1091/mbc.E16-07-0552.

Meier, S.M. et al. (2023) “Multivalency ensures persistence of a +TIP body at specialized microtubule ends,” Nature Cell Biology, 25(1), pp. 56–67. Available at: 10.1038/s41556-022-01035-2.

Meier, S.M., Steinmetz, M.O. and Barral, Y. (2024) “Microtubule specialization by +TIP networks: from mechanisms to functional implications,” Trends in Biochemical Sciences. Elsevier Ltd, pp. 318–332. Available at: 10.1016/j.tibs.2024.01.005.

Meziane, M., Genthial, R. and Vogel, J. (2021) “Kar9 symmetry breaking alone is insufficient to ensure spindle alignment,” Scientific Reports, 11(1). Available at: 10.1038/s41598-021-83136-w.

Miesch, J. et al. (2023) “Phase separation of +TIP networks regulates microtubule dynamics,” Proceedings of the National Academy of Sciences of the United States of America, 120(35). Available at: 10.1073/pnas.2301457120.

Miller, R.K. et al. (2006) “The CLIP-170 Orthologue Bik1p and Positioning the Mitotic Spindle in Yeast,” Current Topics in Developmental Biology, pp. 49–87. Available at: 10.1016/S0070-2153(06)76002-1.

Miller, R.K., Matheos, D. and Rose, M.D. (1999) The Cortical Localization of the Microtubule Orientation Protein, Kar9p, Is Dependent upon Actin and Proteins Required for Polarization, The Journal of Cell Biology. Available at: http://www.jcb.org.

Miller, R.K. and Rose, M.D. (1998) Kar9p Is a Novel Cortical Protein Required for Cytoplasmic Microtubule Orientation in Yeast, The Journal of Cell Biology. Available at: http://www.jcb.org.

Molodtsov, M.I. et al. (2016) “A Force-Induced Directional Switch of a Molecular Motor Enables Parallel Microtubule Bundle Formation,” Cell, 167(2), pp. 539–552.e14. Available at: 10.1016/j.cell.2016.09.029.

Moore, J.K., Silva, S.D.’ and Miller, R.K. (2006) “The CLIP-170 Homologue Bik1p Promotes the Phosphorylation and Asymmetric Localization of Kar9p □ D,” Molecular Biology of the Cell, 17, pp. 178–191. Available at: 10.1091/mbc.E05-06.

Morelli, K. et al. (2025) “A fluid droplet harvests the force generated by shrinking microtubules in living cells,” Newton, 1(8). Available at: 10.1016/j.newton.2025.100197.

Olieric, N. et al. (2010) Automated seamless DNA co-transformation cloning with direct expression vectors applying positive or negative insert selection. Available at: http://www.biomedcentral.com/1472-6750/10/56.

Pereira, G. et al. (2000) The Bub2p Spindle Checkpoint Links Nuclear Migration with Mitotic Exit a kinase with a role in SPB duplication and checkpoint control (Winey et al and Mad3p are part of, Molecular Cell.

Pereira, G. et al. (2001) Modes of spindle pole body inheritance and segregation of the Bfa1p-Bub2p checkpoint protein complex, EMBO Journal.

Shin, Y. and Brangwynne, C.P. (2017) “Liquid phase condensation in cell physiology and disease,” Science. American Association for the Advancement of Science. Available at: 10.1126/science.aaf4382.

Song, X. et al. (2023) “Phase separation of EB1 guides microtubule plus-end dynamics,” Nature Cell Biology, 25(1), pp. 79–91. Available at: 10.1038/s41556-022-01033-4.

Stangier, M.M. et al. (2018) “Structure-Function Relationship of the Bik1-Bim1 Complex,” Structure, 26(4), pp. 607–618.e4. Available at: 10.1016/j.str.2018.03.003.

Wiegand, T. et al. (2025) “Actin polymerization counteracts prewetting of N-WASP on supported lipid bilayers.” Available at: 10.1073/pnas.

Woodruff, J.B. et al. (2017) “The Centrosome Is a Selective Condensate that Nucleates Microtubules by Concentrating Tubulin,” Cell, 169(6), pp. 1066–1077.e10. Available at: 10.1016/j.cell.2017.05.028.

Yaguchi, K. and Woodruff, J.B. (2026) “Functions and regulated material states of cytoskeletal condensates,” Current Opinion in Cell Biology, 100. Available at: 10.1016/j.ceb.2026.102628.

Yin, H. et al. (2000) Myosin V orientates the mitotic spindle in yeast. Available at: www.nature.com.

