## Supplementary Figures for "Low-affinity binding motif in microtubule plus-end condensates specializes microtubule function"

### Supplementary Figure 1

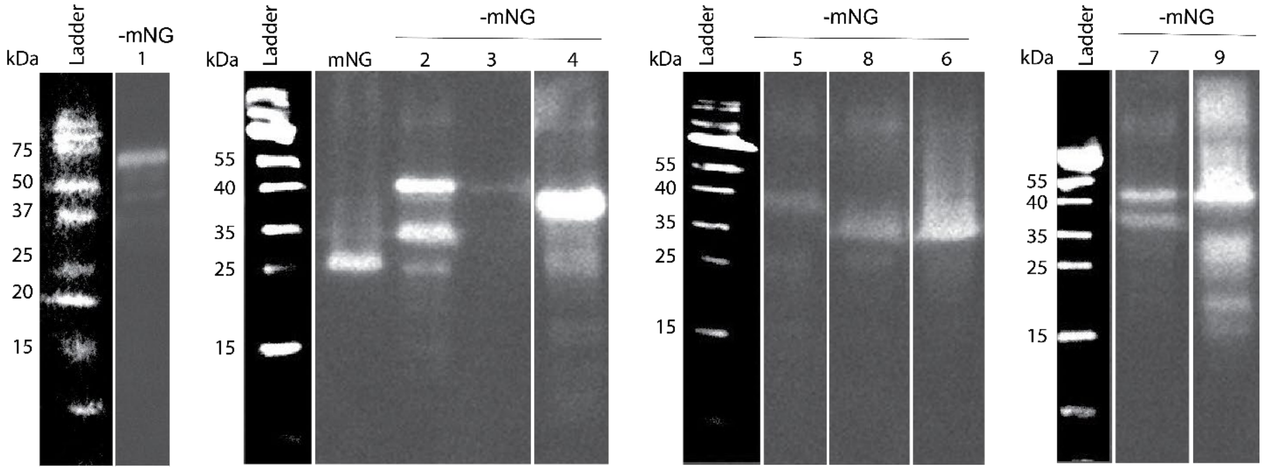


**Figure S1:** Western Blot detection of all N-terminally 6xHis tagged Kar9 fragments with Anti-His. Each panel represents 1 single experiment, with white lines representing un-relevant lanes which have been removed from the figure. Molecular weights of the fragments are as follows: 1: 57.4 kDa, 2: 42.89 kDa, 3: 43.71 kDa, 4: 36.05 kDa, 5:36.00 kDa, 6: 36.57 kDa, 7: 36.30 kDa, 8: 35.94 kDa, 9: 37.42 kDa, mNG: 29.16 kDa.

### Supplementary Figure 2

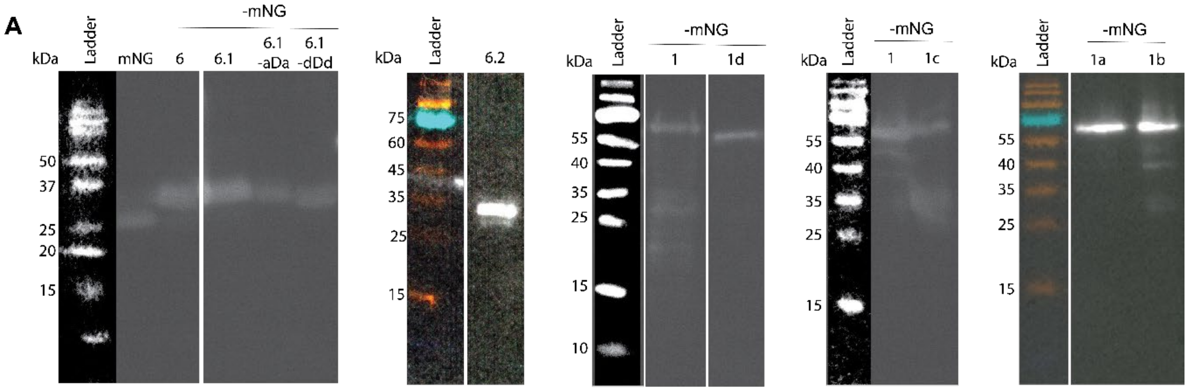


**B** **Manual alignment of interacting motifs in Kar9 homologs**

```
S. cerevisiae   ETPLMAKNKSVLDIEKDKWNHYRSLPSRIPIYK 579
S. cerevisiae   KWNHYRSLPSRIPIYKDKVVKVTVENTPIAKVF 596
S. paradoxus    ETPLLAKNKSVLDVEKDKWRHYQSLPSKIPTYK 579
S. paradoxus    KWRHYQSLPSKIPTYKDKSMKVAVENTPIAKIF 596
S. boulardii    ETPLMAKNKSVLDIEKDKWNHYRSLPSRIPIYK 579
S. boulardii    KWNHYRSLPSRIPIYKDKVVKVTVENTPIAKVF 596
S. uvarum       QTPLMAKNKSVLDIEKDKWKHYQSRPSRIPIYK 581
S. uvarum       KWKHYQSRPSRIPIYKDKPTKFTVENTPVGKVL 598
S. eubayanus    QTPLMAKNKSVLDIEKDKWKHYQSRPSRIPIYK 580
S. eubayanus    KWKHYQSRPSRIPIYKDRPTKLTVENKLAGKVL 597
E. gossypii     QTSLRDVLKTRIQMFRDRDIINLNYLMRLLNHK 197
E. gossypii     VAEVTNTLADKWLVLREKVDPILPKETESDPID 326
Z. rouxii            *MLCFMERLAKDKKRYYDLVGIINKIEP 27
Z. rouxii       SKRTIIHIDHLTKLLKEKHKEVLDKYNIMVKEV 215
N. castellii    LLKKINENDDLAQTIKDRFNSQLAKKSKIITKT 281
N. castellii    KIKFYASKKSLIPSSKEKTSKTIIEQSLTSISS 531
N. dairenensis  NKIDVGVQNYLIASRKEKLSHTMDNSKEPFGLW 663
C. glabrata     ARRDMTNIDSLAEFLRDKYKVLMKKYEFMSSEI 290
C. glabrata     KKLNSVEEIGINVQIREKMSHQMTQKSLTIERT 357
```

**Figure S2A:** Western Blot detection of N-terminally 6xHis tagged Kar9 fragments with Anti-His. Each panel represents 1 single experiment, with white lines representing un-relevant lanes which have been removed from the figure. Molecular weights of the fragments are as follows: mNG:29.16 kDa, 6: 36.57 kDa, 6.1: 33.06 kDa, 6.1-aDa: 33.03 kDa, 6.1-dDd: 32.94 kDa, 6.2: 32.00 kDa, 1: 57.4 kDa, 1a: 57.3 kDa, 1b: 57.3 kDa, 1c: 57.2 kDa, 1d: 56.69 kDa. **S2B:** Manual alignment of KDK motifs of Kar9 homologs across fungal species. Grey-shaded residues represent conserved amino acids. Hydrophobic amino acids (L/I/V) are represented in green while S/T are represented in blue. KDK motifs and amino acids that are part of the motifs are represented in black and bold.

43 **Supplementary Figure 3**

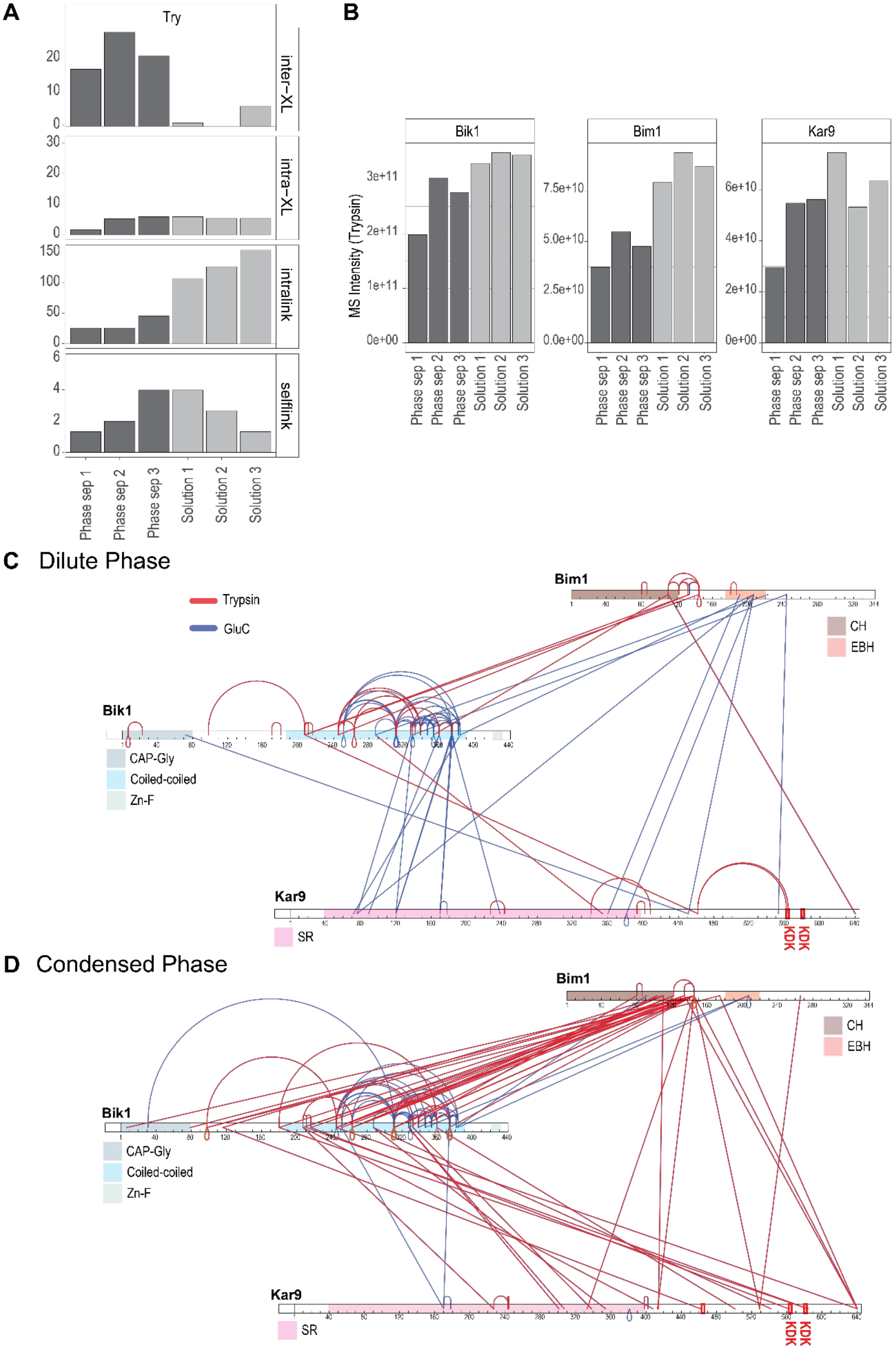

**Figure S3A:** Number of crosslinked identified in the crosslinking dataset in triplicates using trypsin as protease. **S3B:** Protein levels for Bim1, Bik1 and Kar9 in the crosslinking dataset in triplicates using trypsin as protease. **S3C, D:** Visualization of cross-linked peptides for the ternary protein assembly formed by Bim1, Bik1 and Kar9 identified under dilute (500mM NaCl) **(S3C)** and condensed (150mM NaCl) **(S3D)** conditions. The cross-linked peptides identified after trypsin proteolysis are annotated in red, and the peptides identified after GluC proteolysis are annotated in blue. The domain abbreviations in the figure are as follows; CH: Calponin Homology domain, EBH: End Binding Homology domain, CAP-Gly: Cytoskeleton-Associated Protein-Glycine-rich domain, Zn-F: Zinc Finger domain, SR: Spectrin Repeat.

**Supplementary Figure 5**

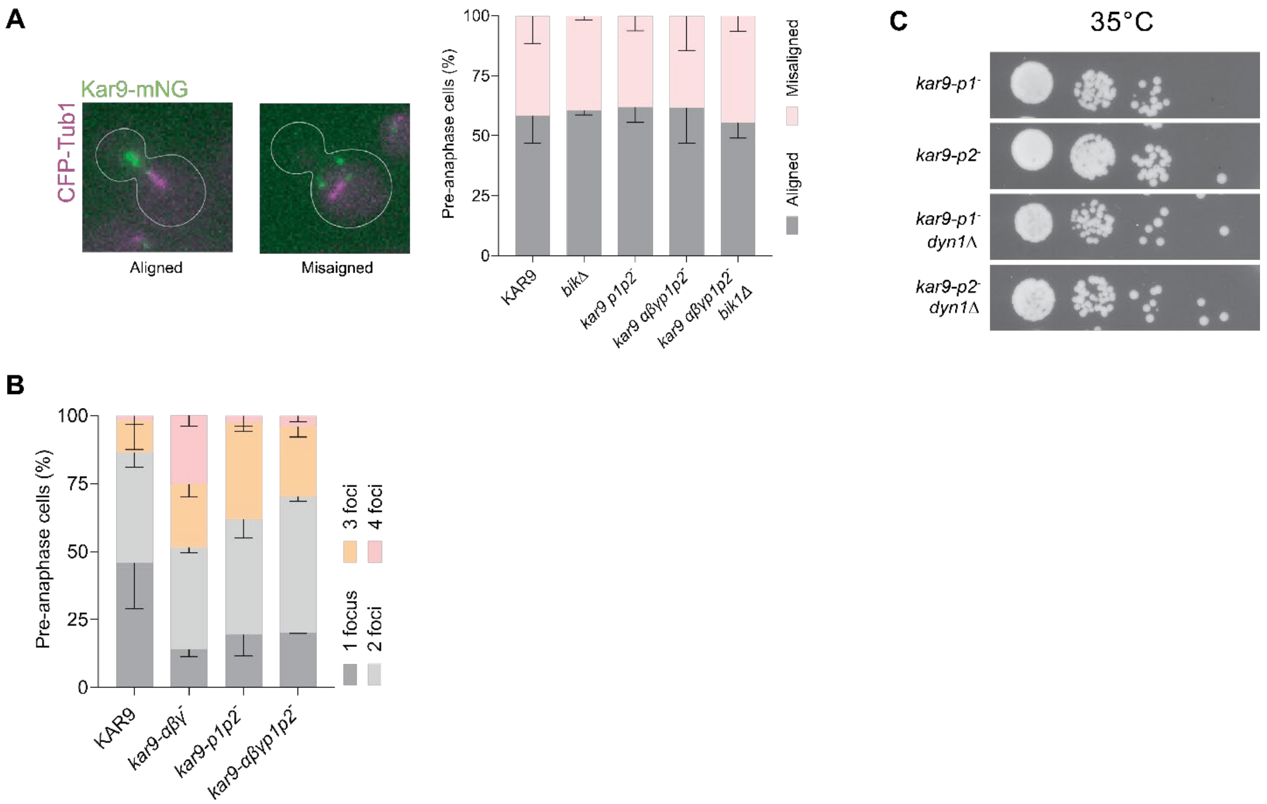


**Figure S5A:** Analysis of alignment of pre-anaphase spindles. **S5B:** Quantification of number of +TIP body foci in different strains in pre-anaphase cells. Error bars represent standard deviations. **S5C:** Spot assay to check viability of control strains harboring different Kar9 variants at 35°C.

### Supplementary Figure 6

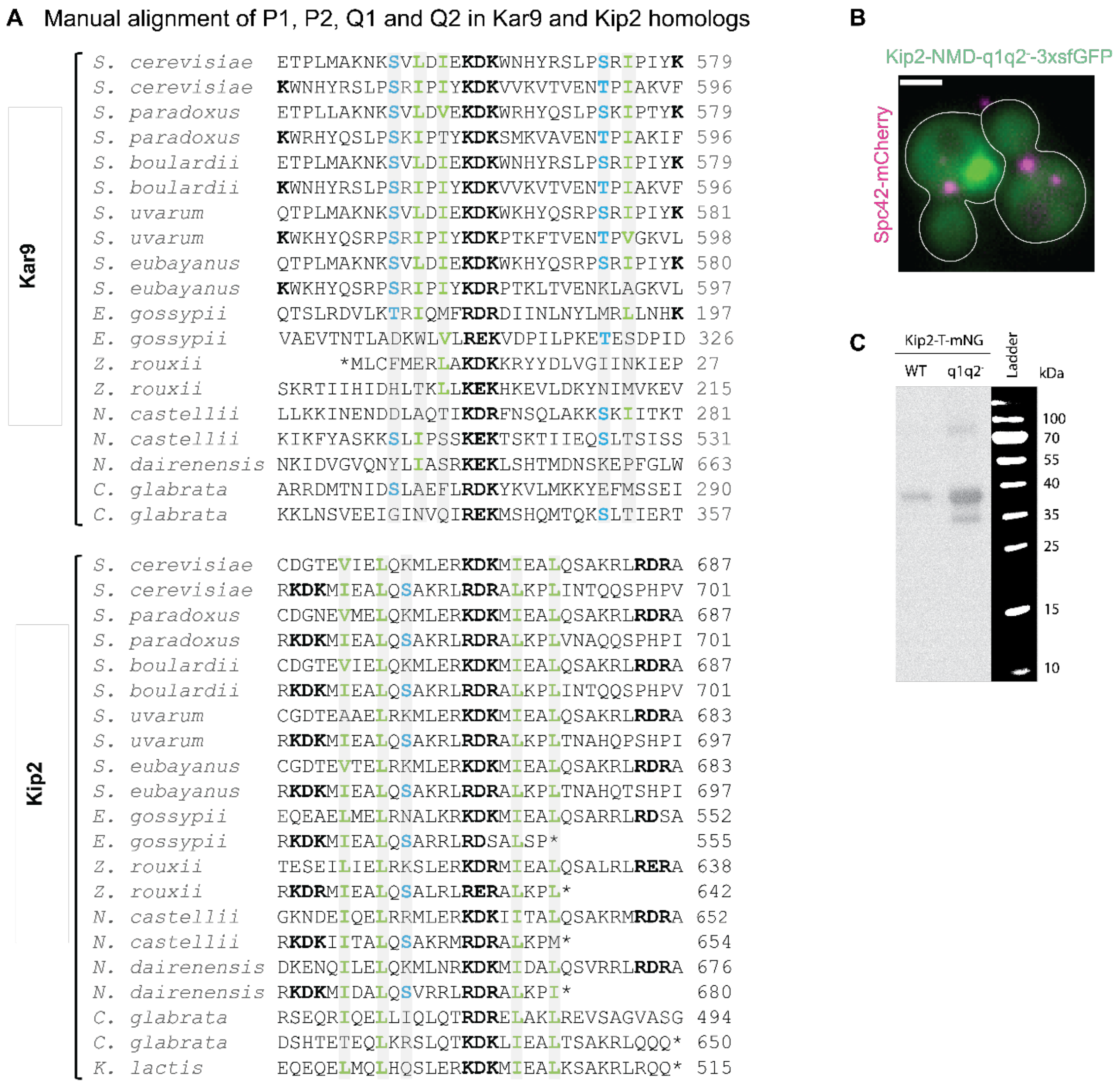


**Figure S6A:** Manual alignment of P1, P2, Q1 and Q2 along with surrounding amino acids in Kar9 and Kip2 homologs. Grey-shaded residues represent conserved amino acids. Hydrophobic amino acids (L/I/V) are represented in green while S/T are represented in blue. P1, P2, Q1, Q2 and amino acids that are part of the motifs are represented in black and bold. **S6B:** Example of a Kip2-NMD-q1q2$^{-}$-3xsfGFP cluster in a metaphase cell (left) with a metaphase cell with unclustered Kip2-NMD-q1q2$^{-}$-3xsfGFP (right). **S6C:** Western Blot detection of all N-terminally 6xHis tagged Kip2-T-mNG fragments with Anti-His. Molecular weights of the fragments are as follows: Kip2-T-mNG: 36.09 kDa, Kip2-T-q1q21$^{-}$-mNG: 35.81 kDa.
